# Neonatal Peyer’s patch cDC activation as a pacemaker of postnatal immune maturation

**DOI:** 10.1101/2022.03.03.482839

**Authors:** N. Torow, R. Li, T. Hitch, C. Mingels, S. al Bounny, N. van Best, E.-L. Stange, A. Benabid, L. Rüttger, M. Gadermayr, S. Runge, N. Treichel, D. Merhof, S. Rosshart, N. Jehmlich, M. von Bergen, F. Heymann, T. Clavel, F. Tacke, H. Lelouard, I. Costa, M. W. Hornef

## Abstract

Marked differences exist between the mucosal immune system of the neonate and adult host. The pronounced influence of the enteric microbiota in adults suggests a causal relationship between postnatal colonization and immune maturation. However, using metagenomic, metaproteomic, and functional immunological analyses we demonstrate an early presence of bacteria and immunogenic microbial antigens preceding immune maturation in the small intestine, the primary inductive site of intestinal immunity. Instead, transcriptomic, flow cytometric and histological analysis indicated neonatal Peyer’s patch (PP) mononuclear phagocytes (MNP) as rate limiting factor of postnatal immune maturation. Despite the early presence of MNPs, conventional dendritic cells (cDC) of type 1, 2a and 2b exhibited significant age-dependent differences in tissue distribution and cellular composition. Single cell transcriptional profiling and functional assays revealed decreased antimicrobial and antigen processing/presentation capacity, an overall retarded cell maturation and reduced antigen uptake. In cDC2a this resulted in a reduced proportion of CCR7^+^ migratory cells and a consequent defect in CD4 T cell priming. Interestingly, transcriptional profiling of neonatal DC subsets identified reduced expression of type I interferon (IFN)-stimulated genes (ISG). Type I IFN induction by oral administration of the TLR7 agonist R848 accelerated MNP maturation and enhanced cognate antigen CD4 T cell priming. However, humoral responses to oral vaccination in the presence of R848 were significantly reduced. Together, our results identify PP MNP maturation as pacemaker of postnatal mucosal immune priming, indicate the biological role of delayed maturation and demonstrate that targeted interventional strategies allow manipulation of mucosal responses in early life.

## Introduction

The neonatal immune system differs significantly and in many aspects from the immune system of the adult individual in both mice and men. These differences are mainly attributed to the need for exposure to environmental stimuli to complete cell differentiation and maturation. Postnatal bacterial colonization and thus exposure to immunogenic microbial antigens and immunostimulatory molecules in the small intestine, the primary site of mucosal immune induction, may determine immune maturation after birth. Likewise, a critical impact on the mucosal immune system has been attributed to the enteric microbiota in the adult host (Gaboriau-Routhiau et al., 2009; Weinstein and Cebra, 1991). Here, microbial signals drive tissue differentiation, stimulate immune maturation and polarization, and reinforce mucosal barrier integrity facilitating a stable ecological niche and maintaining host-microbial homeostasis (Abdel-Gadir et al., 2019). The postnatal period is of particular importance since it exerts a non-redundant and long-lasting influence on various aspects of mucosal immunity (Cahenzli et al., 2013; Constantinides et al., 2019; Gollwitzer et al., 2014; Henrick et al., 2021; Olszak et al., 2012; Ozkul et al., 2020). Reduced microbial exposure during this early time window has been associated with an adverse outcome in various inflammatory and immune-mediated disease models in later life (Cahenzli et al., 2013; Lynn et al., 2018; Olszak et al., 2012; Zanvit et al., 2015). Also in humans, early life alterations in microbial exposure and gut microbiota composition as well as antibiotic use have been associated with an increased risk for allergic and immune-mediated diseases (Kirjavainen et al., 2019; Schuijs et al., 2015; Vatanen et al., 2016b).

However, neonates and adults also differ in their immunological and metabolic needs. Therefore, whereas some differences might result from the requirement for exogenous signals, others might be due to developmental regulation and reflect the specific adaptation of the neonate to its particular situation after birth such as for example the need to maintain energy homeostasis, prevent inappropriate immune activation and sustain tissue integrity and function (Ganeshan et al., 2019; Harbeson et al., 2018; Kuma et al., 2004). Here, small intestinal Peyer’s patches (PP), the main anatomical site of early T cell homing in the murine small intestinal mucosa, are of particular interest (Torow et al., 2015a). As mucosal inductive sites PP are instrumental to intestinal immune homeostasis (Bunker et al., 2015; Pabst and Slack, 2020). PP lack afferent lymphatics but instead sample antigen directly from the small intestinal lumen. In the adult host, an array of conventional and monocyte-derived dendritic cells organized in an intricate microarchitecture positioned between subepithelial dome (SED) and interfollicular regions (IFR) then prime T cells (Luciani et al., 2021).

Despite the recognition of a timed succession of non-redundant phases of immune maturation and priming during postnatal immune development, the factors that determine the postnatal pace of mucosal immune maturation have remained ill-defined prompting our investigation (Herzenberg and Herzenberg, 1989; Hornef and Torow, 2020). Unexpectedly, we found an adult-like bacterial density and substantial antigen diversity and immunogenicity of the enteric microbiota in the murine small intestine early after birth making reduced global antigenicity of the early microbiota an unlikely pacemaker of intestinal immune priming. Instead, we observed delayed anatomical distribution and retarded maturation trajectories of antigen presenting cells (APC) in PP. This maturation blockade was associated with reduced expression of interferon (IFN) stimulated genes (ISG) and type I interferon induction partially reversed this block and enhanced T cell priming. On the other hand, IFN stimulation adversely affected vaccination-induced humoral responses. Our results suggest that APC functionality represents a rate-limiting step for the establishment of intestinal immune homeostasis. A better understanding of immune priming in early life may help to unravel the functional importance of this critical time window for immune homeostasis and ultimately allow manipulation of immune effector function in the neonatal host.

## Results

### Lack of adaptive immune maturation in the neonatal small intestine despite substantial bacterial density and antigenicity

In the murine host thymic output is initiated around birth and naïve T cells populate the lymphoid tissues thereafter. In contrast to adult animals, CD4 T cell numbers were unaffected by the absence of the microbiota in mice at postnatal day (PND)11 (Figure 1A). Despite early homing, the percentage of CD44^hi^ CD4 T cells increased only > 4 weeks after birth but remained low throughout the postnatal period (Figure 1B). To test how quickly an adaptive immune response can be mounted within the neonatal PP environment we transferred naive CD44^lo^ OTII cells from adult mice to 11-day-old mice and measured their activation after oral administration of a high dose of ovalbumin (OVA). The majority of OTII cells had upregulated CD44 expression 48 h after OVA administration in contrast to a much lower percentage of CD44^hi^ cells among endogenous CD4 T cells in neonatal PP illustrating that an immune response to a cognate antigen can be mounted in the neonatal patch environment (Figure S1A). Delayed postnatal bacterial colonization and exposure to immunogenic antigens might therefore explain the slow pace of mucosal immune maturation after birth.

**Figure 1.**
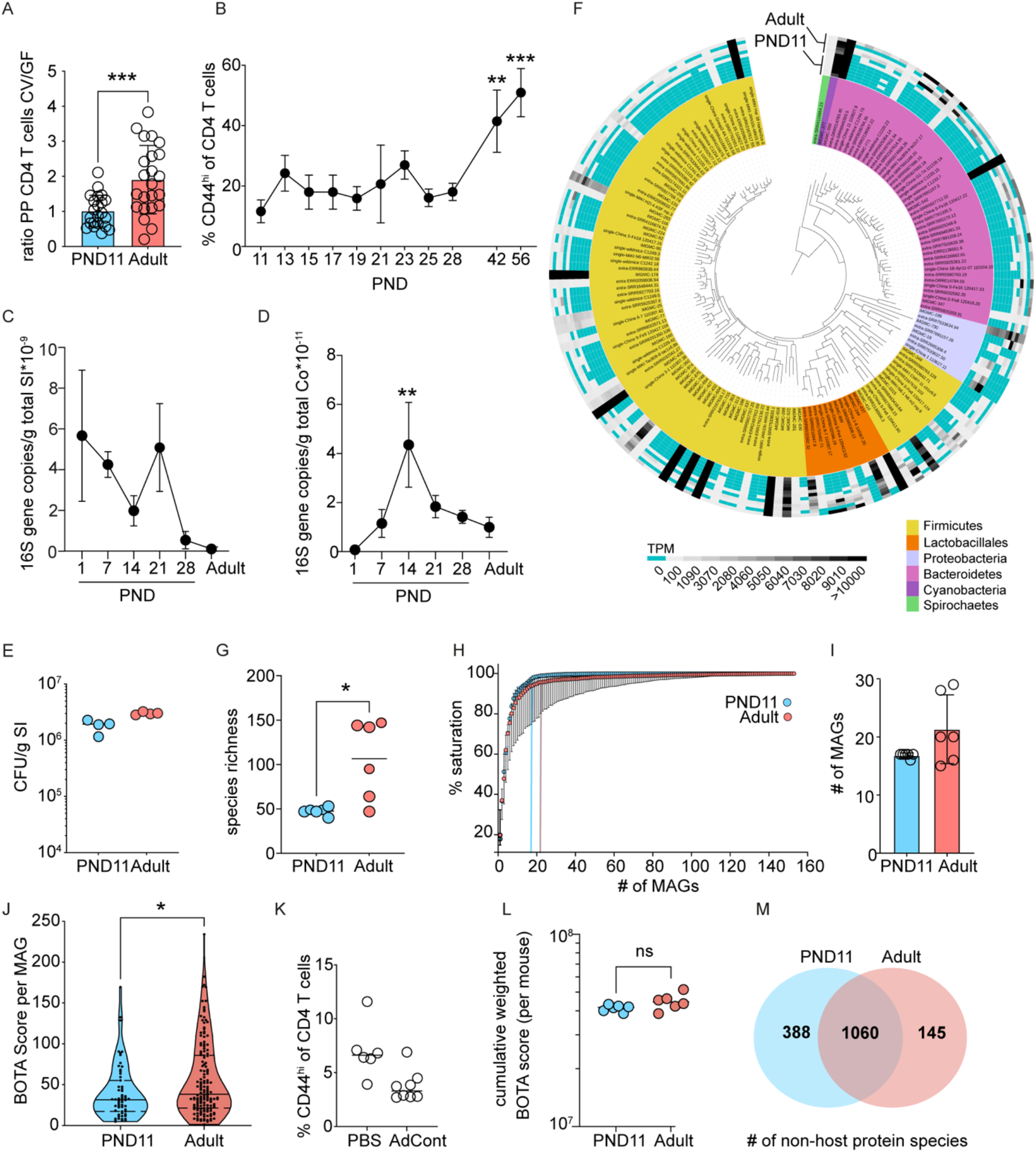
Rapid postnatal establishment of a dense and immunogenic small intestinal core microbiome in the absence of adaptive immune maturation. (A) Number of CD4^+^ T cells per PP in postnatal day (PND) 11 (blue) and adult (red) special pathogen free (SPF) and germ free (GF) mice (n=22-23 PP, mean+SD, Mann-Whitney U test). (B) Percentage of CD44^hi^ cells among total PP CD4^+^ T cells at the indicated age (n=4-20, mean+SD, 1-way-ANOVA and Kruskal-Wallis test; statistical significance indicated relative to PND11). (C and D) Bacterial density determined by quantitative 16S rDNA qRT-PCR in total small intestinal (C) and colonic (D) tissue homogenate at the indicated time points after birth (n=5, mean+SD, Kruskall-Wallis with Dunn’s multiple comparison to adult) (E) Anaerobic culture of homogenized small intestinal tissue from PND11 and adult mice (n= 4, mean, Mann-Whitney-U test). (F) Phylogenetic tree and phylum assignment (inner circle) and abundance (outer circle) as well as (G) species richness (n=6, mean, Mann-Whitney-U test) of the metagenome assembled genomes (MAG) in the luminal content of PND11 (blue) and adult (red) mice obtained by metagenomic sequencing. (H) Saturation plot depicting the number of MAG *vs*. percentage of MAGs required to represent the entire microbiome (in %) at PND11 (blue) and adulthood (red) (n=6, median+SD). (H and I) Knee points of the individual saturation curves indicated in (H) as vertical lines of the median and in (I) per individual PND11 (blue) and adult (red) animal (n=6, mean+SD, Mann-Whitney-U test). (J) Violin plots of BOTA scores of all MAGs identified by metagenomic sequencing in the lumen of the small intestine of PND11 and adult mice (n_PND11_=59, n_Adult_=149, violin plot, Mann-Whitney-U test). (K) Percentage of CD44^hi^ cells among total PP CD4+ T cells in 14-day-old mice following daily oral gavage of adult intestinal content (AdCont) or PBS on PND1-9 (n=6-8; median). (L) Cumulative abundance-weighted BOTA score from all MAGs identified by metagenomic sequencing in the small intestinal lumen of PND11 and adult mice (n=6, mean, Mann-Whitney-U test). (M) Venn diagram of non-host-derived protein species identified by mass spectrometry in small intestinal luminal material of PND11 (blue) and adult (red) mice (n=6). ns, not significant; *, p<0.05; ** p<0.01; *** p<0.001; ****, p<0.0001.

Unexpectedly, analysis of the postnatal establishment of the microbiota in the small intestine, the primary site of mucosal immune induction, by quantitative PCR for bacterial 16S rRNA genes revealed a high bacterial load as early as 24h post parturition in small intestinal tissue with no further increase between postnatal day (PND) 7 and 56 (Figure 1C). Notably, a different scenario with a delayed increase in the bacterial load was detected in the large intestine (Figure 1D). Similarly, anaerobic culture of luminal material on nutrient-rich agar plates revealed a comparable number of colony-forming units (CFU) per gram of small intestinal tissue in neonates (PND11) and adult mice suggesting a similar overall bacterial load (Figure 1E). Next, we employed metagenomic sequencing of DNA prepared from luminal material of neonatal (PND11) and adult small intestines to comparatively analyze the microbiota composition. Sequence reads were matched against the integrated mouse gut metagenome catalog (iMGMC) (Lesker et al., 2020) and 164 metagenome assembled genomes (MAG) could be identified (Figure 1F, Table S1). The overall species richness was significantly higher in adult than neonatal mice (Figure 1G), but this increase was largely confined to low abundant species, whereas the core microbiome represented by highly abundant MAGs, mostly of the Firmicutes and Bacteroidetes phyla was present early in life (Figure 1F). The calculation of the knee points in the saturation plots confirmed that the number of highly abundant species that led to the relative saturation was not significantly different between both age groups (Figures 1H and 1I).

Finally, differences in the diversity of peptides derived from microbiota that colonizes the neonatal *versus* adult small intestine might explain the observed delay in immune maturation ^1^after birth. Using the MAGs identified in neonate and adult small intestine in combination with a previously reported *in silico* pipeline bacteria-origin T cell antigen (BOTA) predictor to determine peptides likely to be presented on host major histocompatibility class (MHC) II molecules derived from proteins encoded in bacterial genomes (Graham et al., 2018). The BOTA score of all individual 164 MAGs identified revealed a moderately but significantly higher mean BOTA score for MAGs present in adult compared to the neonatal consortium (Figure 1J). To test whether the adult antigenic repertoire would induce T cell priming in the neonate, we gavaged adult intestinal content to neonatal mice daily form birth on until PND9 and analyzed the proportion of CD4^+^ T^EM^ at PND14.Surprisingly, no increase in the proportion of CD4^+^ T^EM^ was detected in PPs of these neonates compared to PBS-treated control mice (Figure 1K). Consistently, the cumulative weighted BOTA score that incorporates the abundance of detected species showed no significant difference between neonate (PND11) and adult mice (Figure 1L). Similarly, proteomic analysis of intestinal content reflecting mainly proteins of highly abundant bacterial taxa revealed a large core proteome of 1060 proteins in both neonatal (PND11) and adult mice and a much smaller number of 388 and 145 age-specific proteins, respectively (Figure 1M, Table S2). Taken together we demonstrate that the murine small intestine is colonized early in life by dominant bacterial taxa, of which the cell density, global composition and antigenicity is overall comparable to that in the adult host. In spite of this, homeostatic adaptive immune maturation is delayed until adulthood.

### Neonatal MNP display reduced antimicrobial and antigen processing capacity

Given the central role of mononuclear phagocytes (MNP) in antigen processing, environmental and microbial signal integration, and T cell priming, their reduced numerical or functional capacity in early life might explain the observed delayed immune maturation. Therefore, we analyzed the anatomical organization and transcriptional profile of PP MNP comparatively in neonatal and adult mice. Initially, we measured the strength and distribution of the pan-phagocyte marker Integrin alpha X (CD11c) normalized to the area of the PP in an age-dependent manner. Interestingly, the total phagocyte density was not different between the neonatal (PND5, PND11), weanling (PND21) and adult PP (Figure 2A, Figure S2A). Next, we subjected MNP from neonatal, weanling and adult mice to flow cytometry (PND11, PND21, adult) and scRNAseq (PND11, adult) aiming to characterize the subset composition and transcriptional profile of the various MNP subsets found in PP (Figures S2B, S2C, 2B and 2C) (Bonnardel et al., 2017; Luciani et al., 2021). Whereas the adult MNP compartment was dominated by cDC2 (SIRPa^+/hi^BST2^-^XCR1^-^ MHCII^+^CD11c^+^) they were significantly reduced in neonatal PP. In contrast, cDC1 (XCR1^+^SIRPa^-^MHCII^+^CD11c^+^) were enriched in neonatal PP. Recently, a cDC2b subset has been described in the spleen and is associated with type 3 immunity (Brown et al., 2019; Papaioannou et al., 2021). This subset (RORgt^+^SIRPa^lo^XCR1^-^MHCII^+^CD11c^+^) was also present in PP and at a significantly higher proportion in early life (Figure 2B). The percentage of monocyte-derived cells (BST2^+^SIRPa^+^XCR1^-^MHCII^+^CD11c^+^) was significantly elevated at PND21 but not PND11 compared to the adult. After quality control scRNA-seq analysis recovered a total of 3659 cells derived from two independent pools of each adult and neonatal PP MNP. The cells clustered in six major clusters that were assigned to either type 1, 2a, and 2b cDC or monocyte-derived cells (MC) (Figure 2C). The age-dependent changes in the MNP subset composition in the scRNAseq analysis confirmed the results obtained by flow cytometry (Figure 2D). cDC1 and cDC2a each formed two separate clusters representing immature or quiescent (q) and migratory or activated (act) DC (Cabeza-Cabrerizo et al., 2021). The maturation process from qDC to actDC was accompanied by substantial transcriptional changes leading to their distinct clustering. It was characterized by a discrete set of marker genes that was largely overlapping between actDC1 and actDC2a indicating functional similarities of activated cDC1 and 2a. In contrast, qDC1 and qDC2a were transcriptionally distinct (Figures 2C and S2C). To show that the cDC2b subset truly corresponded to conventional DCs and not type 3 innate lymphoid cells (ILC3) we compared the six clusters to the gene expression profiles of cells available on ImmGen (Figure 2F). Indeed, the cDC2b subset transcriptional profile showed the greatest similarity to the reported intestinal *lamina propria* cDC2 profile. Comparative analysis of the transcriptional profiles of the four MNP subsets in PND11 and adult mice revealed that genes involved in (I) antigen presentation and processing (e.g. *Cd74, H2-Eb1, H2-Dmb2, H2-Aa, H2-Ab1, H2-K1*), (II) microbicidal activity (IFN) signaling (e.g. *Ifitm1, Ifitm2, Ifitm3*) were significantly downregulated in neonatal cells of a given subset compared to their adult counterparts (Figure 2E). Interestingly, actDC1 and actDC2a were transcriptionally very similar between the different age groups (Figure S2D). Neonatal MNP were enriched in transcripts associated with stemness (*Lgals1 and Birc5*). We could confirm the altered expression profile of CD74, H2-K1, and lysozyme on protein level using FACS

**Figure 2:**
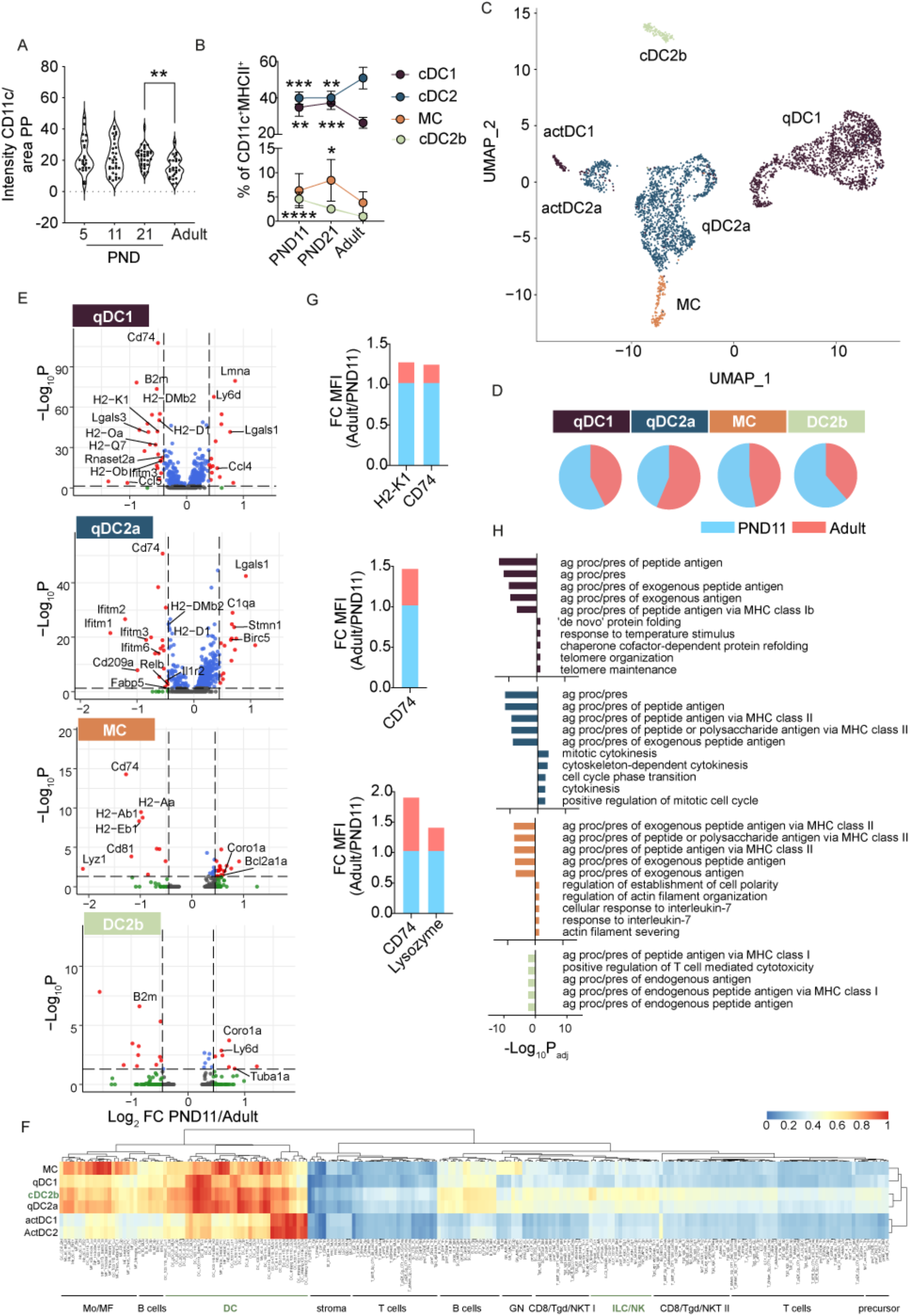
Neonatal PP MNP exhibit diminished antimicrobial activity and reduced antigen processing and presentation capacity. (A) Violin plot of the CD11c positive area normalized to the total PP area at the indicated postnatal age (PND) or in adult mice (n=25-36 follicles; 1-way-ANOVA, Kruskall-Wallis test). (B) Percentage of XCR^+^Sirpa^-^ cDC1 (brown), SIRPa^+^BST2^-^XCR^-^ cDC2 (blue), BST2^+^SIRPa^+^XCR1^-^MC (orange), and RORgt^+^SIRPa^lo^XCR1^-^BST2^-^ cDC2b (gray) among CD11c^+^MHCII^+^CD45^+^live phagocytes isolated from PP of PND11, PND21 and adult mice quantified by flow cytometry (n=5-12; mean+SD; 1-way-ANOVA, Kruskall-Wallis test). (C) UMAP of the scRNAseq analysis of 3659 FACS-sorted CD11c^+^MHCII^+^CD45^+^IgA-TCRb^-^ cells isolated from PP of PND11 and adult mice (n=2; cells pooled from one litter (PND11) or 3 animals (adult) per sample, 6-8 PP per mouse). (D) Pie chart showing the relative contribution of PND11 (blue) and adult (red) cells among the indicated MNP subsets as defined in (C). (E) Volcano plots depicting differentially expressed (DE) genes in CCR7^-^ cDC1, CCR7^-^ cDC2a, monocyte derived cells (MC), and cDC2b (top to bottom panel, subsets as defined in (C)) between PND11 and adult mice. (F) We compared pseudobulk profiles of scRNA-seq clusters from (C) against all ImmGen gene expression profiles using a cosine similarity metric as in (Wang et al., 2021). Rows are normalized by z-score. cDC2b and the ImmGen DC and ILC clusters are highlighted in green. (G) Fold change (FC) of the mean fluorescence intensity (MFI) of the indicated surface proteins between PP cDC1, cDC2, and MC of PND11 (blue, set as 1) and adult (red) mice (n=pooled litter/4 adult animals). (H) Top 5 GO terms under- and overrepresented in the transcriptomic analysis of PND11 cells of the indicated MNP subset (top to bottom: cDC1, brown; cDC2, blue; lysoMNP, orange; cDC2b, gray). ns, not significant; *, p<0.05; **, p<0.01; ***, p<0.001; ****, p<0.0001.

(Figure 2G). Consistently, gene ontology (GO) enrichment analysis confirmed that genes associated with antigen processing and presentation were significantly underrepresented in neonatal cells, whereas cell division and morphology associated pathways were overrepresented compared to adult mice (Figure 2H). Together, the results demonstrated a significant overrepresentation of cDC1 and cDC2b and an overall reduced expression of genes associated with the function of PP MNP in innate defense and antigen processing/presentation in neonatal mice.

### Altered anatomical localization of MNPs in neonatal PP

We next asked whether the overall anatomical PP distribution or the T cell to phagocyte ratio were different in early life. Both would influence the stochastic probability of T cells to recognize their cognate antigen and could explain the observed lack of homeostatic T cell activation. Quantitative flow cytometric analysis of the ratio of antigen-presenting MHCII^+^CD11c^+^ cells and CD4 T cells in PP did not reveal any difference between PND11 and adult mice (Figure 3A). Also, the overall MNP tissue distribution did not reveal any significant age-dependent phenotype. Radial segmentation of the CD11c signal detected by immunofluorescence in adult PP revealed a CD11c signal enriched towards the outer edges of the PP that accommodate the subepithelial dome (SED) and interfollicular region (IFR), as well as the serosal side of the follicle. Enrichment towards the outer PP edges led to a negative slope of the line displaying radial (circumferential to central) signal intensity (Figure 3B). The mean slope was neutral at PND5 indicating a somewhat more equal distribution of CD11c^+^ PP cells during the first week of life but became and remained negative from PND11 on, suggesting an overall adult-like anatomical distribution of CD11c^+^PP cells from PND11 onwards (Figure 3C). However, MNP subsets display a sophisticated anatomical organization in the adult PP (Da Silva et al., 2017). In the adult, cDC1 are mostly found in the IFR. In contrast, cDC2a migrate from the outer edges of the SED to the IFR accompanied by a maturation process that is marked by down-regulation of CCR6, and up-regulation of CD11b and then CCR7 (Bonnardel et al., 2017). Monocyte derived phagocytes, especially lysoDC, are highly enriched at the top of the FAE where they are closely associated with the epithelium and retrieve particulate antigen (e.g. microbial cells). Upon stimulation, they, too, can upregulate CCR7 and migrate to the IFR, but at a much lower rate (Wagner et al., 2020). We therefore used confocal spectral immunofluorescence microscopy to dissect the microarchitecture of the different MNP subtypes. Like in the adult animal, TLR3^+^IRF8^+^CD11c^+^CX3CR1^-^ cDC1 localized to the IFR also in the neonatal PP (Figure 3D). In contrast, marked differences in the MNP distribution between neonatal (PND11) and adult mice were found in the SED and FAE (Figure 3E). SIRPα ^+^CD11c^+^CX3CR^-^Lysozyme^-^ cDC2 appeared more abundant in the FAE of neonatal than adult mice. In neonatal mice (PND11) they were also not restricted to the base of the SED but present in high numbers at the top of the SED interacting intensely with the epithelium (Figures 3E and S3A). Inversely, Sirpa^+^CD11c^+^ CX3CR1^+^Lysozyme^+^CD4^-^ lysoDC were more dominant in the adult SED. This resulted in a significantly altered cDC2:lysoDC ratio with an approximately two-fold enrichment of cDC2 over lysoDC in the neonatal PP (Figure 3G). We also for the first time visualized cDC2b characterized by the expression of the transcription factor RORγ t. Interestingly, RORγ t^+^CD11c^+^CX3CR1^-^CD4^-^ cDC2b were readily detected in the PP of neonatal mice. They localized both to the SED and IFR suggesting a similar maturation/migration pattern as for cDC2a (Figure 3F). Taken together, our results demonstrate that despite a similar MNP to CD4+ T cell ratio and overall distribution, differences in the tissue localization of MNP subsets exist in the neonate consistent with altered cell maturation and function.

**Figure 3:**
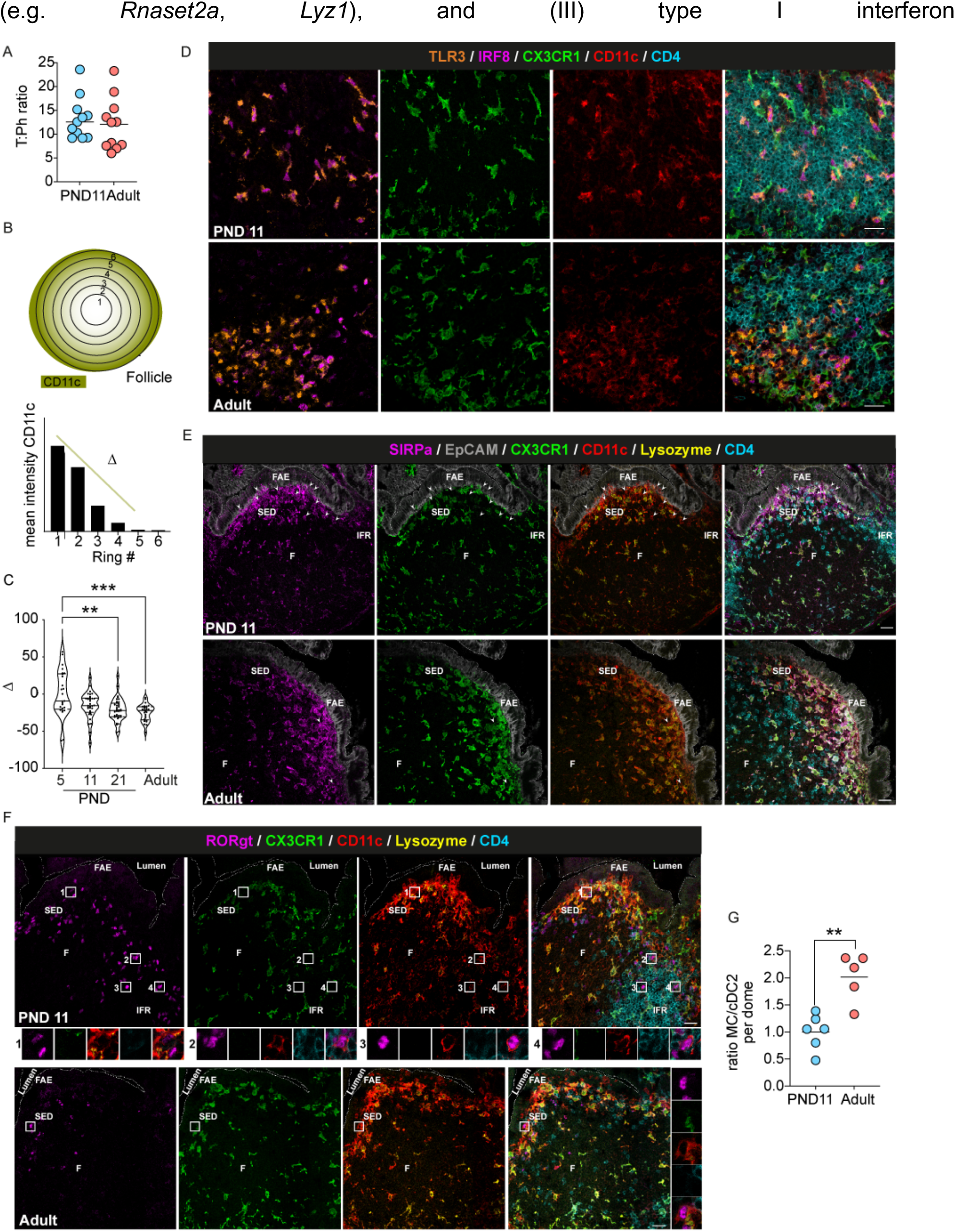
Anatomical distribution of MNP in neonatal and adult PP. (A) Ratio between CD4^+^TCRb^+^CD45^+^live T cells and MHCII^+^CD11c^+^CD45^+^live phagocytic cells in the PP of PND11 and adult mice determined by FACS. (B) Signal distribution model of CD11c+ cells using radial segmentation of the PP area applied to obtain Δ (C) Violin plot of Δ calculated in individual PP of the indicated age (n=25-36 follicles, Violin plot, Kruskall-Wallis test). (D-F) Spectral confocal imaging projection representative of PND11 and adult PP from CX_3_CR1-GFP^+/-^ mice. Sections illustrate the anatomical distribution of (D) cDC1 using the markers TLR3, orange; IRF8, magenta; GFP, green; CD11c, red; CD4, blue (note that only TLR3 IRF8 double positive cells represent cDC1) of (E) cDC2 using the markers SIRPa, magenta; EpCAM, grey; GFP, green; CD11c, red; lysozyme, yellow; CD4, blue (arrowheads point towards cDC2 in the SED) of (F) cDC2b using the markers RORγ t, magenta; GFP, green; CD11c, red; lysozyme, yellow; CD4, blue (boxed areas of individual cDC2b are shown below). Bars, 20 µ m; FAE, follicle associated epithelium; SED, subepithelial dome; F, follicle; IFR, interfollicular region. (G) Ratio between SIRPa^+^CX3CR1^+^Lysozyme^+^CD11c^+^ MC and SIRPa^+^CX3CR1^-^Lysozyme^-^CD11c^+^ cDC2 in the SED quantified in PP as shown in (E) (n=5-6, median, Mann-Whitney-U test. Each dot represents one animal (A) or one follicle (C, G). **, p<0.01; ***, p<0.001; ****, p<0.0001.

### Age-dependent changes of the cDC maturation trajectory

Since changes in the anatomical localization of cDC have recently been associated with a stepwise maturation process (Bonnardel et al., 2017) we next analyzed the maturation profile of MNP subtypes during the postnatal period. We utilized clustering and generated pseudotime trajectories to retrace the developmental paths of quiescent cDC1, cDC2a, and total cDC2b and assessed the conversion rate into the final maturation state of DC marked by sharp changes in their transcriptional profile (Figure 2C) and upregulation of the chemokine receptor CCR7 (Forster et al., 2008). Interestingly, the relative contribution of neonatal cells to the pseudotime trajectory of quiescent cDC1 steadily decreased with advancing degree of maturation as defined by decreasing S-score indicating a strong developmental delay in the maturation of quiescent cDC1 in neonatal PP (Figures 4 A and S4A). However, the proportion of CCR7^+^ migratory cDC1, albeit low in general, was not altered in PND11 mice compared to their adult counterparts (Figures 4A and 4B). In contrast, pseudotime analysis was unable to identify maturational changes in neonatal quiescent cDC2a but suggested a decreased conversion rate into CCR7^+^ actDC (Figure 4D). Sub-clustering of the cDC2a from Figure 2C identified the marker genes *Birc5, Ccr6, Plet1, Itgam* (gene encoding for CD11b), and *Ccr7* that represented cDC2a cells at different maturation stages (Figure 4E). This allowed to study cell maturation by flow cytometry utilizing Plet1 (maturation state (M) 1), CD11b (M2), and CCR7 (M3) as markers for the last three successive developmental stages in the life cycle of cDC2a (Figure 4F). Comparing those cDC2a developmental stages between PND11 and adult PP cDC2a, we detected marked differences with an increased proportion of cells in the M1 state but decreased levels of M2 and M3 in the neonate (Figure 4G). Again, this indicated a skewing towards a maturational delay of neonatal cDC2. Pseudotime analysis revealed an early maturational delay of neonatal cells also for DC2b, but further subdivision of this subtype was limited due to the low total cell number (Figure 4I).

**Figure 4:**
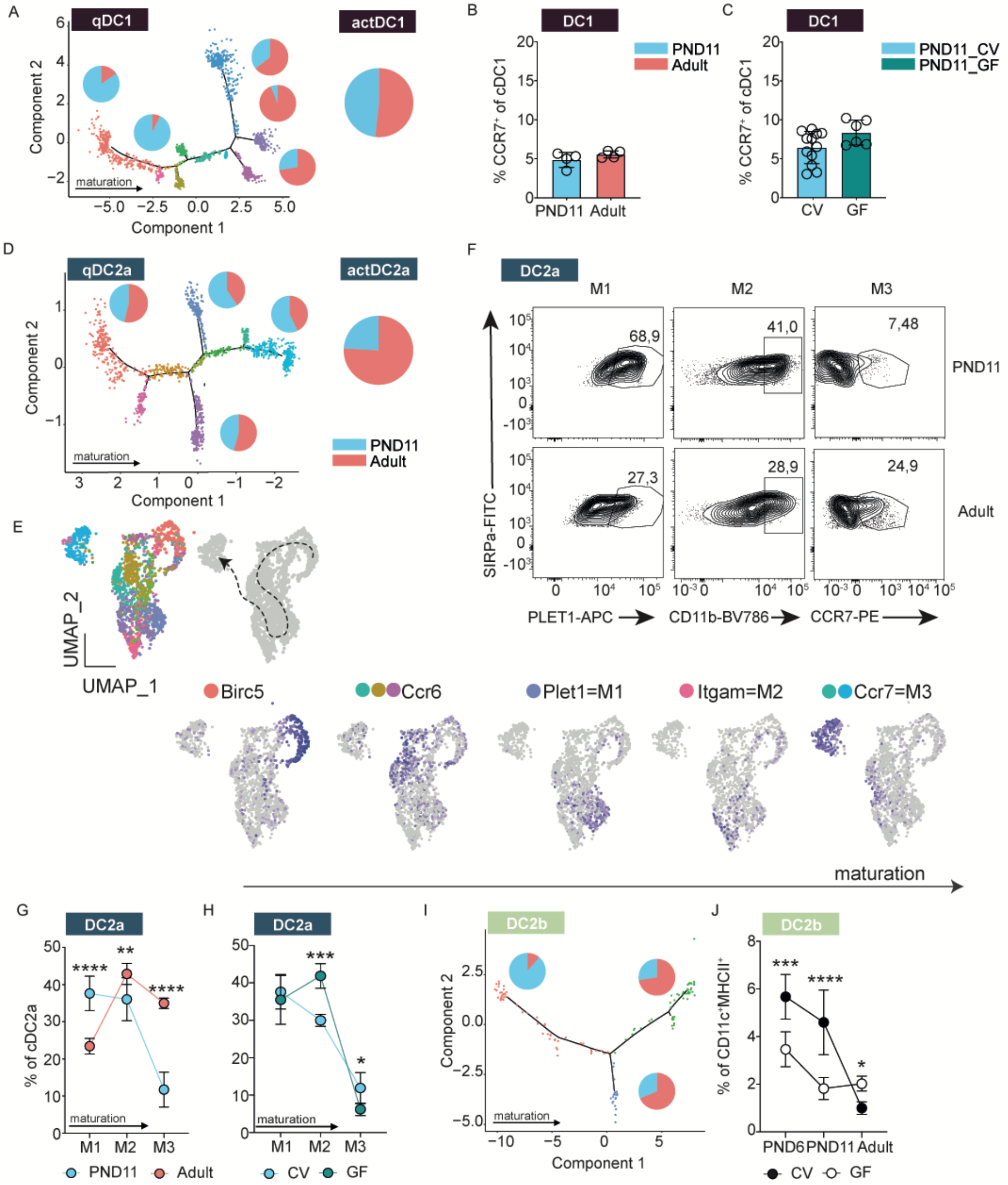
Maturation delay of PP cDC subsets during the postnatal period. (A) Monocle pseudotime trajectory of quiescent CCR7^-^ cDC1 (qDC1) with relative contribution of adult (red) and PND11 (blue) cells to the indicated branches (left panel) and relative contribution of adult (red) and neonatal (blue) cells to the migratory CCR7^+^ cDC1 (actDC1) subset (right panel). (B and C) Percentage of CCR7^+^ cells among XCR1^+^SIRPa^-^MHCII^+^CD11c^+^CD45^+^live cDC1 in (B) conventional PND11 (blue) vs. conventional adult (red) animals (n=4, mean) and (C) PND11 conventional (CV) vs. PND11 germ free (GF) animals (n=6-12, mean) determined by FACS. (D) Monocle pseudotime trajectory of quiescent CCR7^-^ cDC2a with relative contribution of adult (red) and PND11 (blue) cells to the indicated branches (left panel) and relative contribution of adult (red) and neonatal (blue) cells to the activated CCR7^+^ cDC2a subset (right panel). (E) Seurat clusters contributing to cDC2a (left panel, as defined in Fig. 2C), UMAPs indicating marker genes for maturational stages such as BIRC5, CCR6, PLET1 (=M1), CD11b (=M2) and CCR7 (=M3) among cDC2a (middle panels) and proposed maturation trajectory (arrow) (right panel). The cluster color code does not correspond to the pseudotime color code. (F) Representative FACS plots of PND11 and adult cDC2 showing the markers PLET1, CD11b, and CCR7 for maturational stages M1, M2, and M3, respectively. (G and H) Maturational stages characterized by Plet1 (=M1), CD11b (=M2) and CCR7 (=M3) as percentage of total SIRPa^+^BST2^-^XCR1^-^MHCII^+^CD11c^+^CD45^+^live cDC2a in (G) PND11 (blue) vs. adult animals (red) (n=8-10, mean+SD, Kruskall-Wallis test, statistical significance between age groups within the same maturational stage) and (H) PND11 germ-free (GF) vs. conventional (CV) animals (n=4-12, mean+SD; Kruskall-Wallis test, statistical significance between colonization groups within the same maturational stage) determined by FACS. (I) Monocle pseudotime trajectory of total cDC2b with relative contribution of adult (red) and PND11 (blue) cells to the indicated branches. (J) Proportion of RORgt^+^SIRPa^lo^XCR1^-^ cDC2b of total MHCII^+^CD45^+^live MNP at the indicated ages in germ-free (GF) *vs*. conventionally raised (CV) animals (n=4-11, mean+SD, 1-way-ANOVA, Kruskall-Wallis test, statistical significance indicated between the same age group). Each dot represents one animal (B-C); ns, not significant; *, p<0.05; **, p<0.01; ***, p<0.001; ****, p<0.0001.

The global small intestinal microbiota evolves rapidly after birth (Figure 1A) but individual microbiota-derived innate immune signals involved in DC maturation might still arise only with time. We next therefore asked whether cues from the establishing microbiota contribute to the observed maturation of cDC and may account for the detected age-related differences. Despite comparable absolute numbers and a similar subset composition the number CCR7^+^ migratory cDC1 was independent of the presence of a viable microbiota (Figures 4C and S4B). In contrast, cDC2 of germ free PND11 mice displayed an altered maturation profile with increased proportion of cells in the M2 state (CD11b^+^) but decreased M3 levels (CCR7^+^) suggesting that microbial signals contributed to the final maturation step of cDC2 from M2 to M3 and thus towards becoming activated DC in the neonate (Figure 4H). Interestingly, we could also observe an enrichment of cDC2 at M2 in animals deficient for the adaptor protein MyD88 in myeloid cells (MyD88^Δ CD11c^), through which a significant part of microbiota-dependent innate immune signaling is conveyed (Hooper et al., 2012) (Figure S4C). Mice colonized with a wild microbiome (“wildlings”) have been shown to better mimic the human immune system and contain higher numbers of antigen-experienced lymphocytes in the adult (Beura et al., 2016; Rosshart et al., 2019; Rosshart et al., 2017). Thus, we investigated whether this maximal homeostatic microbial exposure impacts PP MNP composition and maturation status in the neonate. We did not observe differences in subset composition or differences at the M2 stage of cDC2a in neonatal PP (Figure S4D). Consistent with recent reports of increased cCD2b numbers after antibiotic administration (Brown et al., 2019), we found an increased proportion of cDC2b in adult germ-free as compared to conventionally raised mice (Figure 4J). In striking contrast, neonatal germ-free mice exhibited a significantly lower proportion of RORγ t^+^ cDC2b than age-matched conventionally bred mice. Together, we identified an early maturation block in cDC1 and cDC2b and a maturation deficit at a later stage in cDC2a in the neonatal PP. Further, microbial signals influenced the maturation of cDC2a and cDC2b.

### Reduced uptake of particulate antigen and CD4^+^ T cell priming in neonatal PP

PP represent the primary site of luminal sampling and processing of particulate antigen and the transcriptomic analysis of neonatal MNP suggested a reduced potential to process and present antigen (Figure 2). Therefore, we next compared the uptake of orally administered fluorescent 100 nm latex beads by SED phagocytes between neonatal and adult PP (Figure 5A). Whereas >90% of sections of adult PP contained bead-positive phagocytes in the subepithelial dome, only approximately 40% of neonatal PP phagocytes had internalized beads at 6 h after oral gavage (Figure 5B). Also, after normalization to the SED surface area, the number of intraphagocytic beads in the PP was significantly reduced in the neonatal host (Figure 5C).

**Figure 5:**
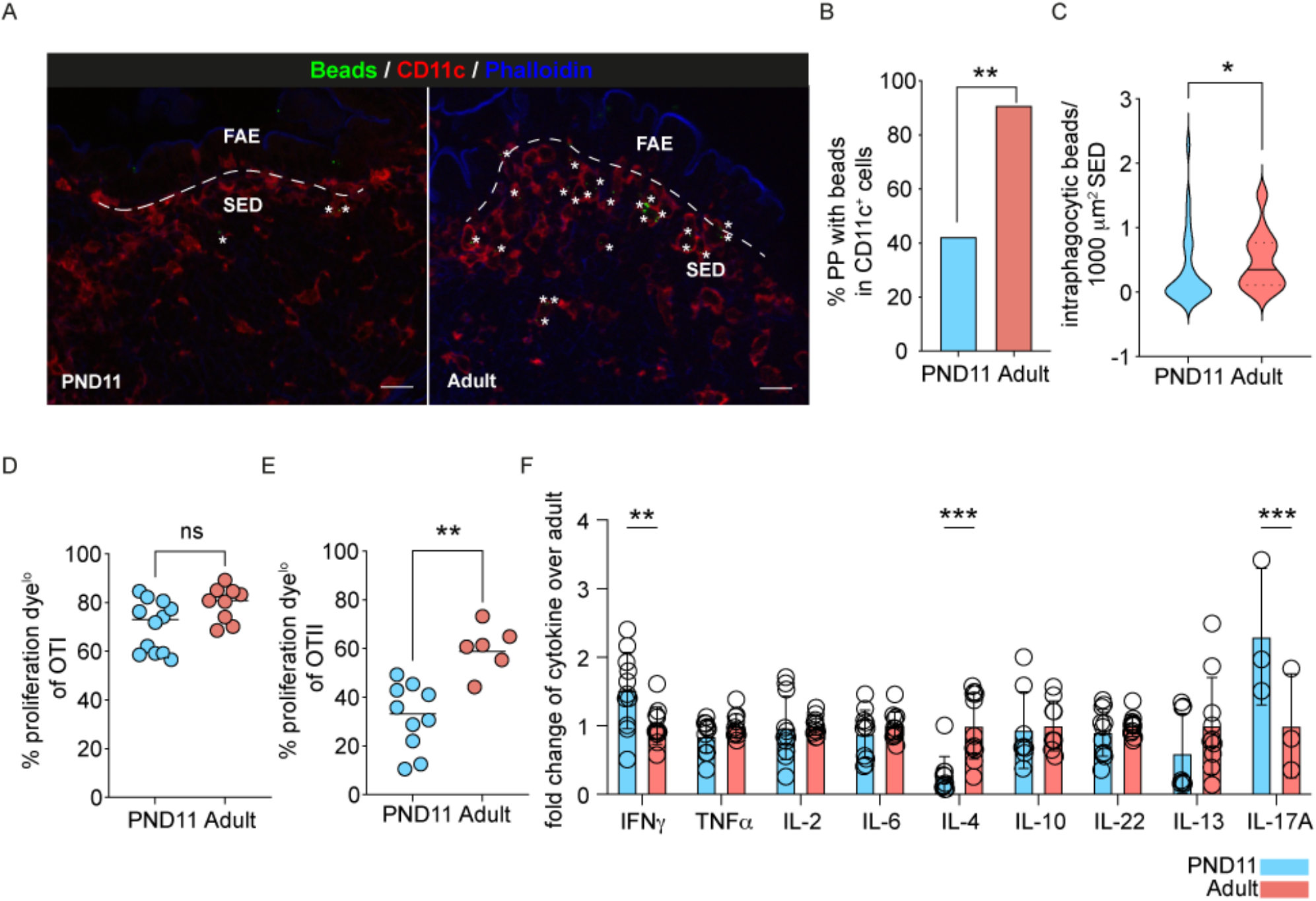
Reduced antigen uptake and altered T cell priming by neonatal PP MNPs. (A) Immunofluorescence imaging representative of PND11 and adult PP at 4h after oral administration of 100 nm fluorescent beads. Sections were stained for CD11c (red), phalloidin (blue), beads are shown in green, asterisks indicate intraphagocytic beads in the SED. Bars, 20µm. (B) Percentage of PP with beads in phagocytic (CD11c^+^) cells within the SED of PND11 (blue) and adult (red) mice (n=23-26 follicles, Fisher’s exact test). (C) Violin plot of intraphagocytic beads normalized to the SED area (n=23-26 follicles, Mann-Whitney U test). (D and E) Proliferation of transferred (D) OTI and (E) OTII cells in PP of PND11 and adult mice 48h after oral administration of OVA determined using the eF670 dilution dye (n=3-11; Mann-Whitney U test). (F) T helper cytokine levels in the supernatant of 5-day co-cultures of OTII cells and MHCII^+^CD11c^+^CD45^+^live MNP isolated from PP of PND11 (blue) and adult (red) mice in the presence of OVA (n=3-11, mean+SD, Mann-Whitney U test). ns, not significant; *, p<0.05; **, p<0.01; ***, p<0.001.

Next, we functionally analyzed neonatal PP cDC1 and cDC2. Whereas CD4 T cell priming is usually attributed to cDC2, cDC1 cross-present extracellular antigens and primarily prime CD8 T cells but can also facilitate CD4 T cell priming (Brown et al., 2019; Cabeza-Cabrerizo et al., 2021). We individually transferred proliferation dye-labelled TCR-transgenic CD4 (OTII) or CD8 (OTI) T cells specific for ovalbumin comparatively into neonatal and adult recipient animals (Figures 5D and 5E). Dilution of the proliferation dye as a measure of T cell activation was assessed 48 h after OVA gavage. Interestingly, efficient CD8 T cell priming was observed both in neonatal and adult PP (Figure 5D). In contrast, CD4 T cell priming was significantly impaired in neonatal animals (Figure 5E). Finally, we wanted to assess the priming capacity of PND11 and adult PP MNP towards different T helper subsets *in vitro*. We therefore co-cultured sort-purified PP MNP with OTII cells and measured secreted T helper cytokines in the supernatant (Figure 5F). Interestingly, OTII cells co-cultured with neonatal PP MNP secreted enhanced levels of IFNγ and IL-17A but decreased levels of IL-4 consistent with our observation of increased cDC1 and cDC2b numbers in neonatal PP specialized in type I and type III immunity, respectively. Together, our results show significantly reduced uptake of particulate material from the gut lumen by neonatal PP phagocytes and decreased conventional MHCII presentation of soluble antigen and subsequent CD4 T cell priming in neonatal mice. In contrast, soluble antigen was cross-presented to neonatal PP CD8 T cells with adult-like efficiency.

### Stimulation of the reduced ISG profile in neonatal cDC accelerates maturation

Interferon-stimulated genes (ISG) such as *Ifitm1, Ifitm2, and Ifitm3* were observed among the top downregulated genes in neonatal cDC2a but also cDC1 (Figure 2E). Tonic microbiota-induced type I IFN signaling was shown to enhance antimicrobial resistance in adult animals (Schaupp et al., 2020). Therefore, increased type I IFN signaling might also promote maturity and functionality of neonatal PP MNP. To test this hypothesis, we employed the TLR7 ligand R848 previously shown to activate small intestinal LP and PP DC in adult animals via release of type I IFN and TNFα by plasmacytoid (p)DC (Bonnardel et al., 2017; Yrlid et al., 2006). Notably, neonatal mice exhibited a high proportion of PP pDC (Figure S6A) and enhanced expression of the activation marker CD86 at 8 h after oral administration of a R848 dose weight adjusted to levels previously administered to adult animals (Figure S6B). To evaluate the influence of IFN I on the transcriptional profile of the MNP subsets, sorted MHCII^+^CD11c^+^ cells from PP of PND11 and adult mice orally treated with R848 or PBS were subjected to scRNAseq analysis. After removal of contaminants, 10756 cells were included in the analysis and attributed to the cDC and MC subsets (Figure 6A). DE gene analysis (PND11 PBS vs. PND11 R848 and adult PBS vs. adult R848) within the qDC1, qCD2a and total cDC2b subset revealed that both adult and neonatal cells displayed a similar ISG profile with *Isg15, Irf7, Ifi27l2a, and Ifitm3* being among the top 10 upregulated genes in each subset upon R848 treatment (Figure 6B). Pseudotime trajectory analysis of qDC1, qDC2a and total cDC2b confirmed a distinct maturation pattern of DCs in R848-treated animals that was driven by an enhanced ISG profile (Figure 6D). Finally, the transition from quiescent to activated DC was quantified. R848 induced an increase in the proportion of activated cDC, however to a lower extent in neonatal than in adult DC (Figure 6C). Interestingly, whereas the mucosal response to orally administered R848 appeared to be less pronounced in the neonate, the levels of proinflammatory cytokines in the serum, including type I IFN, were markedly higher in neonatal animals as compared to their adult counterparts (Figure 6E). Next, we addressed how orally administered R848 would affect adaptive immune priming and memory formation in neonates. OTII priming in neonatal PP was diminished compared to adult PP (Figure 5E) but oral administration of R848 at 4h after OVA partly reversed this phenotype (Figure 6F). The T_FH_ and mucosal antibody response and memory formation was studied using a vaccination protocol with peracetic acid (PAA)-inactivated, orally administered *Salmonella* Typhimurium (STm) previously described in adult animals (Moor et al., 2016). Unexpectedly, we observed an abrogated anti-STm IgA response in the small intestinal mucosa when R848 was co-administered with the STm vaccine (Figure 6G). The systemic anti-STm IgG and IgM antibody response showed a similar trend (Figures S6C and S6D). Together, we demonstrate a potent response in neonates upon R848 treatment that was able to overcome the maturational arrest of neonatal DC in respect to T cell priming. However, type I IFN stimulation abrogated the humoral response to luminal antigens.

**Figure 6.**
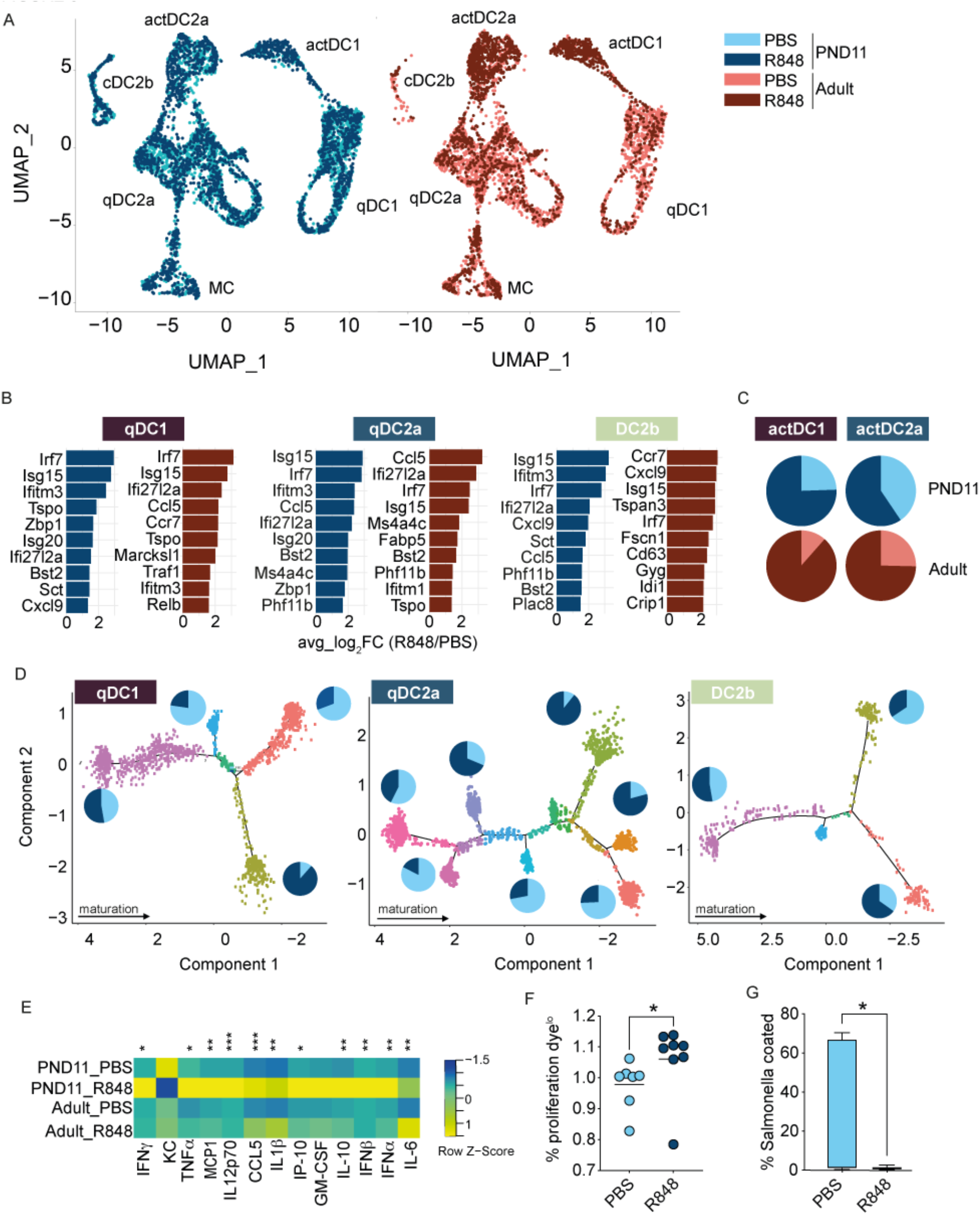
IFN I induces DC maturation in neonatal PP and modifies the adaptive immune response. (A) UMAP of the scRNAseq analysis of 10756 FACS-sorted CD11c^+^MHCII^+^CD45^+^CD19^-^TCRb^-^ cells isolated from PP of PND11 (blue, left panel) and adult mice (red, right panel) orally gavaged with 0.4µg/g bw R848 (dark color) or PBS (light color) (n=1; cells pooled from 4 animals per sample). (B) Top 10 upregulated genes in neonatal (blue) and adult (berry) R848 vs. PBS quiescent DC1, quiescent DC2a and total DC2b. (C) Normalized contribution of activated cDC1 (left) and cDC2a (right) to the activated cluster withing the age group depicted in (A) for PND (blue) or adult (red) cells. (D) Pseudotime trajectory analysis of qDC1, qDC2a and total DC2b in the PND11 group, pie charts represent the relative contribution of cells from PBS (light) and R848 treated animals (dark) to the indicated branches. The branch with the highest S-score was set as the seed of the trajectory. (E) Heatmap of serum cytokine levels measured 8h after oral gavage of 0.4µg/g bw R848 or PBS in PND11 and adult mice (n=4-5; Kruskall Wallis test, comparison within age group, stars indicate significance in PND11 PBS vs. R848). (F) Ratio of proliferation dye diluted OTII cells in PP of neonatal mice 48h after oral gavage of OVA±R848 (n=7-8; Mann-Whitney U test). (G) Percentage of anti-IgA^+^ *Salmonella* incubated with small intestinal washes from adult animals vaccinated with *Salmonella*±R848 as neonates (n=11-12; Box plot, Mann-Whitney U test). Each dot in F and G represents one animal; ns, not significant; *, p<0.05; **, p<0.01; ***, p<0.001.

## Discussion

Despite the differences between mice and men the ontogeny of immune maturation is thought to be of major conceptual relevance. Epidemiological studies in both, humans and mice have identified the early postnatal period as a critical and non-redundant time window for the establishment of a fully functional mucosal immune system (Cahenzli et al., 2013; Constantinides et al., 2019; Depner et al., 2020; Ege et al., 2011; Martinez et al., 2018; Olszak et al., 2012; Ozkul et al., 2020; Vatanen et al., 2016a). Mature T cells have been recently described in the fetal *lamina propria* in humans. However, their TCR specificity is unclear and they might exert tissue remodeling rather than antimicrobial activity (Li et al., 2019; Mishra et al., 2021; Schreurs et al., 2019). The characterization of the pace of postnatal immune maturation and the functional analysis of the adaptive changes in either host are expected to advance our understanding of the mechanisms that facilitate life-long immune homeostasis or determine the susceptibility to inflammatory and immune-mediated diseases.

Our results show that delayed postnatal immune maturation is accompanied by retarded PP cDC differentiation and activation. The potential of type I IFN or IFN-stimulated gene (ISG) products to accelerate cDC activation and enhance immune reactivity in neonates suggests a functional role. Whereas enhanced IFN and ISG expression during viral infection has long been described (Mesev et al., 2019), a number of reports also described tonic expression in healthy animals with comparable levels in primary and secondary immune organs and small intestine (Abt et al., 2012; Ganal et al., 2012; Lienenklaus et al., 2009; Marie et al., 2021; Schaupp et al., 2020). Plasmacytoid (p)DCs represent the most likely source of this tonic type I IFN (Schaupp et al., 2020) but cDCs also directly sense IFN-inducing microbial stimuli (Cabeza-Cabrerizo et al., 2021). Beside its antiviral activity, type I IFN has been noted to enhance mononuclear phagocyte function and adaptive immunity (Gough et al., 2012; Taniguchi and Takaoka, 2001). Type I IFN was shown to alter the transcriptional, epigenetic, and metabolic basal state of cDCs (Schaupp et al., 2020) leading to accelerated maturation (Koya et al., 2017), augmented endocytic and proteolytic activity and enhanced cross-presentation capacity (Date et al., 2020; Schaupp et al., 2020; Spadaro et al., 2012; Zietara et al., 2009).

Since antibiotic treatment or the use of germ-free mice led to reduced ISG expression and an impaired antiviral host response in adult animals (Abt et al., 2012; Ganal et al., 2012; Yang et al., 2021), microbial stimuli have been suggested to drive tonic type IFN expression (Kawashima et al., 2013) and influences homeostatic immune activation in adult and aged mice (Schaupp et al., 2020; Stebegg et al., 2020). This also includes the IFN gene expression profile in DC (Stefan et al., 2020). Interestingly, some bacterial taxa might harbor a particularly strong ISG inducing activity such as lactobacilli, *Blautia coccidioides* and the anaerobic commensal bacteria of the *Bacteroides* species (Ayala, 2021; Kawashima et al., 2018; Kawashima et al., 2013; Stefan et al., 2020; Yang et al., 2021). In our dataset, *Bacteroides spp*. were absent from the PND11 microbiota in our metagenomic analysis and, although found only at low abundance in adult mice, could drive intestinal innate immune stimulation and type I IFN expression (Stefan et al., 2020). Alternatively, infections with unknown viral pathogens or subclinical reactivation of latent or endogenous viruses may contribute to tonic IFN expression.

Unexpectedly, we found that the enteric microbiota of the small intestine exhibits high bacterial density, significant microbial diversity and antigenicity early after birth (Pantoja-Feliciano et al., 2013; van Best et al., 2020). Previous reports described a delayed postnatal increase in bacterial diversity over 2-3 years in humans and 6-8 weeks in mice (Jiang et al., 2001; Roswall et al., 2021; van Best et al., 2020; Yatsunenko et al., 2012). However, these studies analyzed fecal samples that reflect the distal intestinal tract and the situation appears to be markedly different in the proximal intestine. Interestingly, an early high bacterial density and adult-like diversity were also reported in the oral cavity (Koren et al., 2021). Importantly, microbial molecules provide the sole source of non-self antigens in exclusively breastfed newborns. We therefore concluded that it is not the absence of a sufficiently dense or globally antigenically diverse microbiota in the neonate that explains the delayed immune cell activation. Notably, our results do not exclude that specific particularly immunogenic and/or immunostimulatory bacterial species that start to colonize the intestine only during adolescence influence postnatal immune maturation as discussed above.

cDCs of wildling mice, i.e. animals raised in the presence of viral, bacterial, fungal and parasitic microorganisms that are also present in their natural habitat still exhibited a delayed time course of immune maturation. It therefore appears that the phenotype of delayed mucosal immune maturation is not restricted to animals housed under SPF conditions and that the IFN I induction driven by bacterial colonization and exposure to viral pathogens might not suffice to accelerate immune maturation. Thus, despite the fact that we were able to compensate for the delay and accelerate cDC differentiation by interventional strong type I IFN induction, type I IFN (alone) might not represent the physiological stimulus driving DC maturation in adolescent animals. A still undefined stimulus acting upstream of ISGs, a combination of stimuli, an altered susceptibility to those stimuli or a developmental program might contribute. Beside cDCs, other factors might also influence the observed time course in the immune maturation after birth. We and others provided the first evidence for an active suppression of premature T lymphocyte maturation in neonates (Koch et al., 2016; Torow et al., 2015b) but the previously identified mechanisms, namely maternal immunoglobulins and neonatal T regulatory cells, only accounted for a minor part of this suppression. Additionally, microfold (M) cells were previously shown to fully mature only after birth consistent with the low expression of the M cell-specific transcription factor Sip-B as well as known markers of M cells such as Ccl9, UEA1 and gp2 during the first week after birth (Kanaya et al., 2012; Zhang et al., 2014). M cells facilitate uptake and translocation of luminal material but also serve as scaffold for certain types of antigen-presenting cells to sample luminal material for subsequent processing and presentation (Lelouard et al., 2012). Interestingly, we observed reduced numbers of lysoDC in the neonate FAE but enhanced numbers cDC2. These neonatal cDC2 interacted with the epithelium and exhibited membrane extensions towards the FAE surface suggesting an active role in antigen uptake. Similar to what has recently been described in the adult intestinal mucosa, these epithelium-associated cDC2 might be less mature and promote T cell hypo-responsiveness contributing to delayed immune maturation (Rivera et al., 2021). Unexpectedly, we noted a reduced humoral response to the oral vaccine when R848 was co-administered in neonatal mice. The underlying reason is currently unclear. Previous reports in adult animals described both enhanced T cell responses but also increased differentiation of plasma cells, isotype switching and humoral immunity by type I IFN (Jego et al., 2003; Le Bon et al., 2001; Longhi et al., 2009). A critical influence on the differentiation of CD4+ T cells to either plasma cell-promoting follicular helper (T_FH_) or type 1 helper (T_H_1) cells might be the time kinetic of type I IFN production (De Giovanni et al., 2020). Although we detected a strong ISG response in cDC1, 2a and 2b upon R848 administration, we lack information about the precise time kinetic of local type I IFN secretion in the PP *in vivo*. Moreover, a recent study found that cohousing of animals with pet-store mice and thus a history of diverse bacterial and viral exposure dampened the humoral response to vaccination (Fiege et al., 2021).

The observed reduction in the humoral response suggests a potential evolutionary benefit of delayed immune maturation. Humoral immunity critically contributes to the protection against encapsulated bacteria that exhibit the highest incidence in neonates and small infants. An alternative explanation might represent energy allocation trade-offs between immunity and growth. The adult individual with filled energy stores mounts an array of finely tuned innate and adaptive host defense mechanisms aiming at pathogen elimination (Harbeson et al., 2018). In contrast, the neonate host with limited metabolic resources but high demand due to body growth and tissue development may favor disease tolerance (Ganeshan et al., 2019; Harbeson et al., 2018; Medzhitov et al., 2012). It thereby also avoids tissue immunopathology due to inappropriate immune stimulation by the sudden and largely unregulated postnatal exposure to microorganisms (Kuma et al., 2004). Additional explanations might be the prevention of microbiota-induced cross-reactive T cells induced by the lack of terminal deoxynucleotidyl transferase expression and absence of N nucleotide addition during V-(D)-J gene recombination early after birth (Gavin and Bevan, 1995), the preference for early establishment of self-tolerance (Gardner et al., 2008) or the host’s attempt to preserve the available T cell pool for relevant antigens present in the adult microbiota and diet to support life-long mucosal homeostasis (Torow et al., 2015a).

A mechanistic understanding of neonate-specific features and their biological relevance as well as the ability to manipulate immune reactivity might ultimately facilitate targeted interventional strategies to prevent or treat diseases of the term and preterm human neonate (Kollmann et al., 2020). Infections represent the main cause of morbidity and mortality in neonates and young infants worldwide and preventive measures such as vaccines could reduce childhood mortality (Costello and Naimy, 2019; Countdown to 2030 Collaboration, 2018; G. B. D. Child Mortality Collaborators, 2016; Kotloff et al., 2013; Levine et al., 2020; MacLennan and Saul, 2014; Mathers et al., 2017; Oppong et al., 2020; Prudden et al., 2020). The use of vaccines in neonates that carry the largest disease burden, however, is less effective calling for strategies to accelerate immune maturation and improve the immune response to the vaccine antigen in particular at mucosal body surfaces accessible to non-parenteral vaccine administration (Kollmann et al., 2020).

## Material and Methods

### Animals

All animal experiments were performed in compliance with the German animal protection law (TierSchG) and approved by the local animal welfare committees, the Landesamt für Natur, Umwelt und Verbraucherschutz, North Rhine Westfalia (84-02.04.2015.A063 and 84-02.04.2016.A207) and Regierungspräsidium Freiburg, Freiburg (X-20/05F). C57BL/6 wild-type (WT), B6.Cg-Tg(TcraTcrb)425Cbn *Rag1tm1Mom*/J (OTII), C57BL/6-Tg(TcraTcrb)1100Mjb/J (OTI), B6.129P2(SJL)-*Myd88*^*tm1Defr*^ Cg-Tg(Itgax-cre)1-1Reiz/J (MyD88^Δ CD11c^), B6.SJL-*Ptprc*^*a*^ *Pepc*^*b*^/BoyJ (CD45.1), and B6.129P2(Cg)-*Cx3cr1*^*tm1Litt*^/J (CX3CR1-GFP) mice were bred locally and held under specific pathogen–free (SPF) or germ-free (GF) conditions at the Institute of Laboratory Animal Science at RWTH Aachen University Hospital. C57BL/6NTac wildling mice were created through inverse germ-free rederivation as previously desribed (Rosshart et al., 2019), housed as well as bred locally at the animal facility of the Medical Center – University of Freiburg, Germany. C57BL/6NTac murine pathogen free (MPF) control mice were originally purchased from Taconic Biosciences, subsequently bred locally and housed under SPF conditions at the animal facility of the Medical Center - University of Freiburg, Germany. Wildlings and C57BL/6NTac MPF mice were age matched for all experiments.

### *In vivo* models

For R848 administration, PND11 and adult mice were administered with 0.4 µg/g body weight (b.w.) R848 by intragastric gavage. For the OTI/OTII transfer, 1-2×10^5^ eF670 marked OTI or OTII cells/g b.w. were transferred into PND11 and adult mice i.p. and OVA was administered 24 h later by intragastric gavage at 2 mg/g b.w. 48 h after OVA administration, the proliferation of the transferred T cells was assessed. In experiments involving R848, R848 was orally gavaged 4-5 h after OVA at 0.4 µg/g b.w. For exposure of neonatal mice to the adult enteric microbiota, small intestine, ceacum, and colon of adult mice were opened longitudinally and flushed with 5 mL sterile PBS per animal for 5 min. under gentle agitation. Tissues were removed and clumps were suspended by vigilant pipetting. Large matter was removed by brief centrifugation at 300g for 30s and the supernatant was collected. Aliquots of adult intestinal content were stored at -20°C for single use and administered daily by intragastric gavage to neonatal mice starting at PND3 until PND10. For the administration of fluorescent beads, 5×10^11^ yellow-green fluorescent 100 nm beads/g b.w. (Fluoresbrite) were administered by intragastric gavage to PND11 and adult mice and PP were examined 6 h later.

### Vaccination against *Salmonella*

The vaccination protocol from Moor et al.(Moor et al., 2016) was adapted. *Salmonella* Typhimurium (STm) (SL1344) was grown in 500 mL at 37°C and 200 rpm to late stationary phase, harvested by centrifugation (3000 xg, 15 min., 4°C) and resuspended at 10^9^-10^10^ CFU/mL in PBS. Bacterial suspensions were incubated in 10 mL aliquots in 50 mL tubes with 0.04% peracetic acid (PAA) (Sigma) for 1 h at RT, centrifuged at 3000 xg for 10 min. and resuspended at 10^11^ particles/ mL. 100 µl of the suspension was plated to verify complete inactivation. Mice were orally gavaged with 4×10^9^ particles PAA STm/g b.w. ± 0.4 µg/b.w. R848 at PND7 and 14 and with PAA STm but no R848 at PND21. Serum and SI wash were collected at the age of 7-12 weeks.

### Isolation of PP cells

For neonatal PP a dissection microscope was used. PP were excised, counted and digested using 0.2 mg/mL Liberase TH/DNAse I for 45 min at 37°C and then dissociated by vigorous pipetting. Myeloid cells were positively enriched using CD11c MicroBeads (Miltenyi). OTI and OTII cells were isolated from lymph nodes and digested using 0.2 mg/mL Liberase TH/DNAse I for 45 min at 37°C and then dissociated by vigorous pipetting. OTI cells were then further enriched using a naïve CD8 T cell isolation kit (Miltenyi).

### Flow cytometry and cell sorting

Single cell suspensions were blocked with anti-mouse CD16/32 (clone 93, Biolegend, 1:200 dilution) and stained with fluorescently labeled antibodies for 20-40 min. at 4°C. Antibodies were used at a 1:200 dilution unless stated otherwise. CCR7 surface staining was incubated at room temperature for 45 min. at a dilution of 1:50 prior to surface staining with the other antibodies. To determine coating of *Salmonella* with specific immunoglobulins from serum and small intestinal washes the protocol published by Moor et al. (Moor et al., 2016) was used. Blood was collected into clotting activating gel tubes (Sarstedt), incubated 15-30 min. at RT and centrifuged at 16000 xg for 15 min.; serum was stored at -20°C until further use. Small intestines were opened longitudinally, suspended in sterile PBS for 5 min., and vortexed. The tissue was removed and the SI wash was centrifuged at 16000 xg for 10 min.; supernatants were stored at -20°C until further use. Serum and SI washes were inactivated at 56°C for 30 min. and centrifuged at 16000 xg for 10 min. *Salmonella* Typhimurium (SL1344) were grown to late stationary phase in liquid medium, pelleted at 3000 xg at 4°C for 10 min., washed in 0.4 µm filtered PBS, and resuspended at 10^7^ CFU/mL in 0.4 µm filtered 3%FCS/PBS. Sera and SI washes serially diluted (1:2) with an initial dilution of 1:3 and undiluted, respectively. 25 µ l of bacteria were incubated with 25 µ l of serum/SI wash for 1 h at 4°C, washed twice with 200 µ L 0.4 µm filtered 3% FCS/PBS, stained with fluorescently labeled antibodies for 1 h at 4°C and washed before acquisition.

Following antibody clones (Biolegend) were used: anti-CD11b (clone M1/70), anti-CD11c, anti-CD4 (clone RM4-5), anti-CD44 (clone IM7), anti-CD45R/B220 (clone RA3-6B2), anti-CD86 (clone GL1), anti-IA/IE (clone M5/114.15.2, 1:400), anti-XCR1 (clone ZET), anti-BST2 (clone 927), anti-CCR7 (clone 4B12), anti-CD8a (clone 53-6.7), anti-CD19 (clone 6D5), anti-CD45 (clone 30-F11), anti-CD45.1 (clone A30), anti-CD45.2 (clone 104), anti-SIRPa (clone P84), anti-CD74 (clone In1/CD74), anti-mouse IgG1 (clone RMG1-1, 1:40), anti-mouse IgA (clone C10-3, 1:50), anti-mouse IgM (clone RMM-1), anti-RORgt (clone B2D), anti-TCRb (clone H57-597), anti-TCRVa2 (clone 20.1). Homemade PLET1 antibody was provided by C.Ruedl and conjugated to AF647. Dead cells were stained with 7-AAD (Biolegend) or ZOMBIE NIR (Biolegend). Countbrite counting beads (Biolegend) were added to the FACS samples to assess the absolute cell numbers. Cells were acquired on an LSR Fortessa (BD) using BD FACS DIVA software. Cell sorting was carried on a FACS ARIA II (BD) equipped with a 100 mm nozzle using BD FACS DIVA software. Final analysis was performed using FlowJo (v. 10, BD).

### DC OTII co-culture and T helper cytokine induction

PP MNP (MHCII^+^CD11c^+^CD45^+^live) from PND11 and adult wild type mice (pooled litter, 4 adult animals per biological replicate) and OTII (CD4^+^live) were FACS sorted and co-cultured at a ratio of 1:5 in IMDM 10% FCS, 1% penicillin and streptomycin (GIBCO), 1% L-glutamine (GIBCO) and 5 nM 2-mercaptoethanol (Carl-Roth) in the presence of 1 mg/mL OVA for 5 days. Supernatants were analyzed using the Legendplex mouse T helper cytokine panel version 3 (Biolegend) according to the manufacturer’s instructions. Independent experiments were pooled after normalization.

### Absolute quantification of bacterial genomes (16S rRNA gene copy numbers)

The 16S rRNA gene copy numbers were determined in total homogenized small intestinal and colonic tissue samples of C57BL/6J wildtype mice collected at 1, 7, 14, 28, and 56 days after birth (n = 6/age group) and described previously (van Best et al., 2020). Total DNA was isolated from approximately 200 mg frozen tissue by repeated bead-beating combined with chemical lysis and column-based purification using the QIAamp DNA Stool Mini Kit (Qiagen, Hilden, Germany) according to the manufacturer’s instructions (Salonen et al., 2010). Subsequently, the PCR primers 5′-CCTACGGGNGGCWGCAG-3′ (16S_341_F) and 5′-GACTACHVGGGTATCTAATCC-3′ (16S_805_R) were used for amplification (Klindworth et al., 2013). The real-time PCR was performed on a MyiQTM System (BioRad, USA) in a reaction-volume of 25 µl with 12.5 µl iQTM SYBR Green (Biorad), 2 µl template DNA and 0.75 µl primers (10 µM). The cycling conditions were 95 °C for 3 min. followed by 35 cycles of 95 °C for 15 s; 55 °C for 20 s and 72 °C for 30 s. Total 16S gene copy numbers were calculated by comparing the CT value to a standard curve with known concentrations of a plasmid encoding the 16S rDNA gene target sequence of *E*.*coli*. Tissues from germ free animals were used as negative controls, in which no specific PCR product was detected.

### Quantification of bacteria by anaerobic culture

Small intestines were opened longitudinally, quickly transferred to Hungate tubes containing anaerobic PBS (0.05% L-cysteine, 0.02% DTT) with ∼10 glass beads and vortexed. To ensure anaerobic conditions, the supernatant was transferred into another Hungate tube containing anaerobic PBS (0.05% L-cysteine, 0.02% DTT). The suspension was plated onto tryptone soya agar plates with sheep blood (Oxoid) under anaerobic conditions N2 89.3%, CO2 6.0%, H2 4.7%) in a chamber (MBRAUN, Garching, Germany). All materials were brought into the anaerobic workstation at least 24h prior to work. CFU were quantified after 24h of incubation at 37 °C.

### Immunofluorescence and confocal microscopy

PP were excised and fixed using Antigenfix for 45 min. at 4°C and incubated in 30% sucrose/PBS until saturated. Then, they were embedded in OCT compound, frozen over liquid nitrogen and stored at -80°C. Sections were cut at 10µm thickness and slides were stored at - 20°C. After permeabilization and unspecific binding site blocking with PBS containing 0.5% saponin, 2% bovine serum albumin, 1% fetal calf serum, and 1% donkey serum for 30 minutes, sections were labeled overnight at 4°C with primary antibodies followed by washing in PBS before incubation for 1 hour at room temperature with secondary antibodies. Sections were then washed and blocked with sera of primary antibody species for 30 minutes, before staining in the same blocking buffer with fluorochrome-coupled antibodies for 1 hour. Slides were mounted in ProLong Gold and observed with a Zeiss LSM 780 confocal microscope using the spectral imaging mode (Lelouard, 2018).

### Quantification of the CD11c signal distribution in PP

In a first instance all images were manually segmented by encircling each follicle individually with a polygon. This polygon mask served, together with the image, as basis for further processing steps. To determine the distribution of the CD11c signal with respect to the distance to the surrounding shape the following steps were performed: First, the image was filtered with a Gaussian kernel (σ =9, filter size: 201×201) to achieve a smoother surface and to account for the sparse distribution of positive cells within the image. Next, mean pixel intensities were computed within rings with a specified minimum and maximum distance from the masks’ contour. This was obtained using morphological erosion with circular structuring elements of different diameters. To measure the occurrence of the CD11c signal with respect to distance to the contour, we extracted these mean values in non-overlapping, consecutive rings within the whole mask as illustrated in Figure 3B. The thickness (diameter) of each ring was set to 5 pixels to achieve a good trade-off between resolution and smoothness. The center circle was not considered for further processing since it can become arbitrarily small leading to inaccurate measurements. With the obtained mean values, we fit a first order polygon and used the slope Δ as a descriptor for each follicle. The descriptor was normalized for the size of the mask by multiplying the slope with the number of rings.

### Single cell RNA sequencing

Pooled cells from a litter of neonatal mice (5-8 animals) or 3 adult mice were used and samples were analyzed in biological duplicates for the PND11 vs. adult PP MNP experiment and as single samples for the R848 stimulation experiment. MACS enriched cells isolated from PP were subjected to FACS sorting as CD11c+MHCII+CD45+live+IgA-TCRb- and further processed using the Chromium Single Cell 3’ v2 kit (10xGenomics) according to the manufacturer’s protocol. After library generation, sequencing was performed on a NextSeq 550 (IZKF genomics facility of the RWTH Aachen University, Aachen, Germany). Sequencing was performed on Illumina NextSeq 550 (paired-ends, 2×75 bp). With default parameters, we used Cell Ranger (version 2.1.1) to align reads to the mouse genome mm10. Following that, we utilized Seurat (v3.1.0) (Satija et al., 2015) to achieve a high-level analysis of the scRNA-seq data. Cells with high mitochondrial (≧ 10%) and ribosomal (≧ 25%) gene content, as well as cells with ≧ 30.000 UMIs having an increased probability to represent doublets and cells with ≧ 30 detected genes were excluded from further analysis. We used Seurat to regress out cell cycle, mitochondrial, ribosomal, and UMI counts and performed a log-normalization of read counts. In a second step contaminant cell types (B cells, Cd19; pDCs, Siglec; villus macrophages; Fcgr1 (the latter only for the data set examining naïve PND11 vs. adult MNP)) were removed. Canonical correlation analysis (CCA) based on the first 15 CCs was used to combine samples from R848 and PBS experiments (Butler et al., 2018). Unsupervised clustering was performed using a shared nearest neighbors graph with a resolution of 0.8 and k = 15. The following parameters were used to build UMAP representations: min dist = 0.3, max.dim = 2, seed.use = 36). To determine cluster specific markers, we implemented the FindMarkers gene function with an adjusted p value of 0.05. Differential gene expression analysis was performed with Seurat using the Wilcoxon test. Next, we used monocle (version 2.12.0)(Trapnell et al., 2014) in the integrated data to reconstruct cell development trajectories. For the reconstruction of cell development trajectories, we employed the 1000 most relevant genes identified by Monocle’s unsupervised feature selection approach (dpFeature). The root of the trajectory was defined choosing the branch with the highest S-score, which also correlated with the expression of stemness-associated genes such as *Birc5*. For the PND11 PBS vs. PND11 R848 and adult PBS vs. adult R848 scRNA-seq data set data integration was performed by anchors identification based on the results of CCA analysis. The first 20 CCs were used for the anchor identification and first 20 PCs from PCA analysis were used for UMAP representation. For clustering resolution was set to 0.5. Raw data were deposited under GEO accession number GSE188714.

### Metagenomic microbiota analysis

DNA for the metagenomic analysis was isolated using a modified protocol according to Godon et al. (Godon et al., 1997). Snap frozen samples were mixed with 600 µl stool DNA stabilizer (Stratec Biomedical), thawed, and transferred into sterile 2-mL screw-cap tubes containing 500 mg 0.1 mm (diameter) silica/zirconia beads. 250 μl 4 M guanidine thiocyanate in 0.1 M Tris (pH 7.5) and 500 μl 5 % N-lauroyl sarcosine in 0.1 M PBS (pH 8.0) were added. Samples were incubated at 70 °C and 700 rpm for 60 min. A FastPrep® instrument (MP Biomedicals) cooled with dry ice was used for cell disruption (3 × 40 s at 6.5 M/s). Subsequently, 15 mg polyvinylpyrrolidone (PVPP) was added and samples were vortexed, followed by 3 min. centrifugation at 15000 xg and 4°C. Approximately 650 µl of the supernatant was transferred into a new 2 mL tube, which was centrifuged for 3 min. at 15000 xg and 4°C. Afterwards, 500 µl of the supernatant were transferred into a new 2 mL tube and 50 µg of RNase was added. After incubation at 37 °C and 700 rpm for 20 min., the gDNA was isolated using the NucleoSpin® gDNA Clean-up Kit from Macherey-Nagel according to the manufacturer’s protocol. DNA was eluted from columns twice using 40 µl elution buffer and the concentration was determined using a NanoDrop® (Thermo Scientific). Samples were stored at -20 °C. Sequencing of the metagenomic DNA was performed as follows: mechanical shearing of 600 ng metagenomic DNA to a size of 200 bp (200 s, peak incident power: 75 W, duty factor: 10%, cycles per burst: 200) was performed using a Covaris M220. Subsequently the libraries were prepared with the NEBNext® Ultra II DNA Library Prep Kit for Illumina® (NEB, USA) according to the manufacturer’s protocol. The PCR enrichment of adaptor-ligated DNA was conducted with 7 cycles and NEBNext® Multiplex Oligos for Illumnina® (NEB, USA) for single end barcoding. Size selection and clean-up of adaptor ligated DNA was conducted using AMPure beads (Beckman Coulter, USA). The quality (TapeStation, Agilent Technologies, USA) and quantity (Quantus, Promega, USA) of the resulting libraries was checked at the IZKF Core Facility (UKA, RWTH Aachen University) prior to sequencing on a NextSeq500 (Illumina, USA) with a NextSeq500 High Output Kit v2.5 (300 Cycles).

For the quality control, the metagenomic sequencing data were first trimmed using Trimmomatic (Bolger et al., 2014) using a sliding window of 5:20, minimum length of 50, trailing of 3, leading of 3 and illumine clipping values of 2:30:10. Host reads were removed using BBmap (Bushnell, 2014) to map the reads against the genome of Mus musculus GRCm38. Alignment was done using a minimum ratio of 0.9, maximum indel of 3, band width ratio of 0.16, band width of 12, minimum hits of 2 with at least 25 consecutive matches using a kmer length of 14. The first 10bp on each read before mapping and regions with an average score below 10 were also trimmed but undone after mapping (untrim).

The host filtered reads of the metagenomic DNA were then assembled using MEGAhit (Li et al., 2015) using a kmer list of 21,27,33,37,43,55,63,77,83,99, with a minimum multiplicity for filtering kmers of 5. PROKKA was used to predict the proteome from the assembled contigs using default options (Seemann, 2014). The predicted proteome was functionally annotated against the KEGG database using Kofamscan (Aramaki et al., 2020).

Host filtered reads to the iMGMC non-redundant collection of metagenome-assembled genomes (MAGs) (Lesker et al., 2020) were aligned using BBsplit (Bushnell, 2014) with a minimum identity of 90% to produce a coverage statistics file which was converted into TPM via the makeTPMfromCovStats.sh script from the iMGMC best-practices repository. The TPM values were used to calculate the minimal number of species that covered the maximal amount of the data based on the knee point, implemented in python using the Kneed module (Satopa, 2011). Only MAGs with a TPM value >100 in >2 samples were retained for further analysis. A phylogenomic tree of the 153 iMGMC-derived MAGs was generated using their predicted proteomes in Phylophlan3 v3.0.60 (Asnicar et al., 2020). Diversity was set as high using the ‘supermatrix_aa.cfg’ config file. The tree was visualized in iTOL (Letunic and Bork, 2019). Raw data were deposited under BioProject accession number PRJNA794356.

### Antigenic peptide prediction from metagenomic data using BOTA

The proteins encoded on the 153 MAGs recovered from iMGMC were predicted using PROKKA v1.14.6 (Seemann, 2014). The genome, predicted proteome and general features were used as input for BOTA v0.1.0 (Graham et al., 2018) using the H-2-IAb model for MHC class II (allele IA) antigen prediction.

### Metaproteomic analysis

For metaproteome analysis, samples were dissolved in 500 µl SDS lysis buffer (0.29 g NaCl, 1M Tris-HCL pH=8, 5M EDTA pH=8, 0.4g SDS in 100 mL water). Protein extraction was done by bead beating (FastPrep-24, MP Biomedicals, Sanra Ana, CA, USA; 5.5 ms, 1 min., 3 cycles) followed by ultra-sonication (UP50H, Hielscher, Teltow, Germany; cycle 0.5, amplitude 60%) and centrifugation (10,000 xg, 10 min.). The protein lysate was loaded on a SDS-gel and run for 10 min. The gel piece was cut, washed and incubated with 25 mM 1,4 dithiothreitol (in 20 mM ammonium bicarbonate) for 1 h and 100 mM iodoacetamide (in 20 mM ammonium bicarbonate) for 30 min., destained, dehydrated and proteolytically cleaved overnight at 37 °C with trypsin (Promega). The proteolytically cleaved peptides were extracted and desalted using ZipTip μC18 tips (Merck Millipore, Darmstadt, Germany). The peptide lysates were re-suspended in 15 µl 0.1% formic acid and analysed by nanoliquid chromatography mass spectrometry (UltiMate 3000 RSLCnano, Dionex, Thermo Fisher Scientific). Mass spectrometric analysis of eluted peptide lysates was performed on a Q Exactive HF mass spectrometer (Thermo Fisher Scientific) coupled with a TriVersa NanoMate (Advion, Ltd., Harlow, UK). Peptide lysates were injected on a trapping column (Acclaim PepMap 100 C18, 3 μm, nanoViper, 75 μm x 2 cm, Thermo Fisher Scientific) with 5 μl/min. using 98% water/2% ACN 0.5% trifluoroacetic acid, and separated on an analytical C18 column (Acclaim PepMap 100 C18, 3 μm, nanoViper, 75 μm x 25 cm, Thermo Fisher Scientific) with a flow rate of 300 nl/min. The mobile phase was 0.1% formic acid in water (A) and 80 % ACN/0.08 % formic acid in water (B). Full MS spectra (350–1,550 *m/z*) were acquired on the Orbitrap at a resolution of 120,000 with an automatic gain control (AGC) target value of 3×10^6^ ions. Data resulting from LC-MS/MS measurements were analyzed with the Proteome Discoverer (v.2.4, Thermo Fischer Scientific) using SEQUEST HT. Protein identification was performed using a self-built reference database downloaded from UniProt (reference numbers: UP000008827.fasta, UP000019116.fasta, UP000007305.fasta and the predicted proteome from the combined metagenomic assembly). Searches were conducted with the following parameters: Trypsin as enzyme specificity and two missed cleavages allowed. A peptide ion tolerance of 10 ppm and an MS/MS tolerance of 0.05 Da. As modifications, oxidation (methionine) and carbamidomethylation (cysteine) were selected. Peptides that scored a q-value >1% based on a decoy database and with a peptide rank of 1, were considered identified. For the differential analysis of the proteomics data the R package DeqMS 1.8.0 (https://doi.org/10.1074/mcp.TIR119.001646) was used. For each group (PND11 and adult), proteins with valid quantification in at least half of the samples (3/6) were included.

### Statistical analysis

The Mann-Whitney U test was used to compare two groups, the Kruskall-Wallis with Dunn’s multiple comparison was used to compare multiple groups and one variable, 2-way-ANOVA with Bonferroni’s multiple comparison test was used to compare multiple groups and two variables. Statistical tests were carried out using Graphpad Prism and p<0.05 was considered significant. The number of biological replicates analyzed and the statistical test used are indicated in the respective figure legends.

## Supplementary Information

**Supplementary Figure 1:**
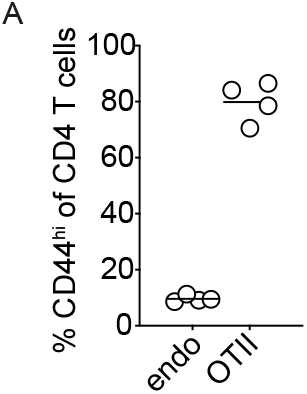
Rapid postnatal establishment of a dense and immunogenic small intestinal core microbiome in the absence of mucosal adaptive immune maturation. **(A)** Percentage of CD44hi cells among endogenous CD4 T cells (endo) and transferred adult OTII cells (OTII) isolated from PP of neonatal PND11 mice 48h after OVA gavage (n=4, mean).

**Supplementary Figure 2:**
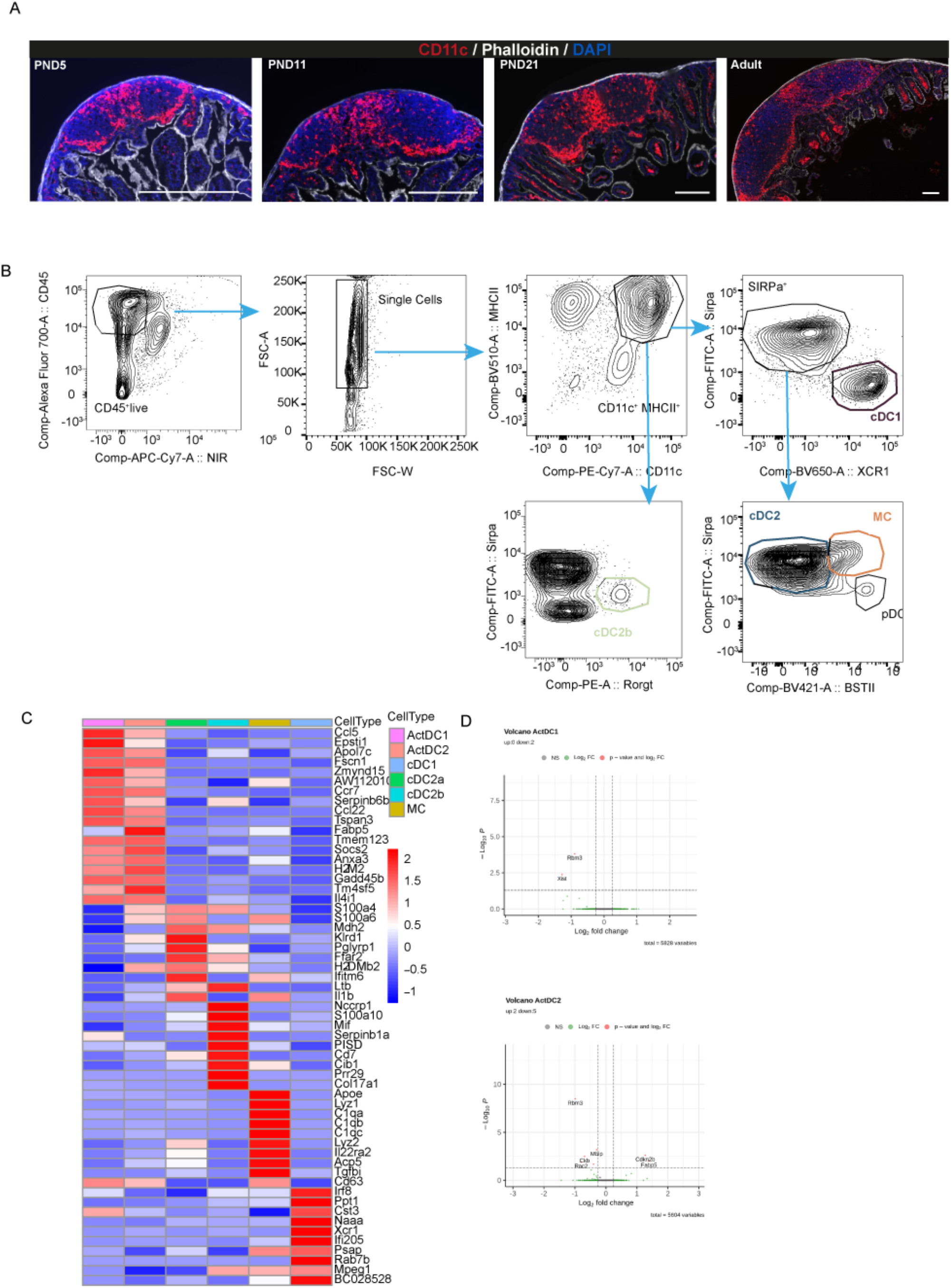
Neonatal PP MNP exhibit diminished antimicrobial activity and reduced antigen processing and presentation capacity. **(A)** Representative immunofluorescent images of PP at indicated ages stained for CD11c (red), phalloidin (white), DAPI (blue) used for quantification of the CD11c signal intensity depicted in Figure 2A; Bars, xyz µm. **(B)** FACS gating strategy to identify MNP subsets in PP. **(C)** Genes defining the cell clusters in the UMAP in Figure 2C. **(D)** Volcano plots of between PND11 and adult mice differentially expressed (DE) genes in actDC1 (upper panel) and actDC2a (lower panel).

**Supplementary Figure 3:**
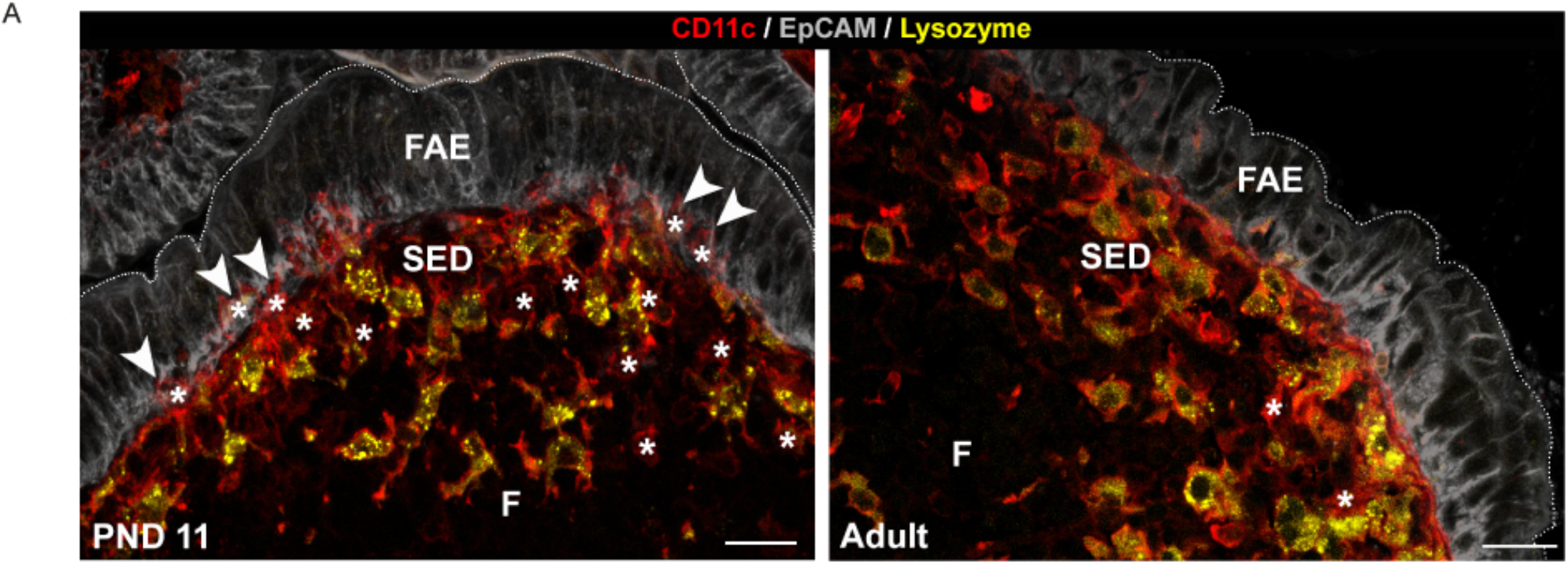
Anatomical distribution of MNP in neonatal and adult PP. **(A)** Spectral confocal imaging projection representative of PND11 and adult PP from CX3CR1-GFP^+/-^ mice. Enlarged image of the FAE/SED area of the 3^rd^ panel in Figure 3E stained for EpCAM, grey; CD11c, red; lysozyme (Lyz), yellow); Bars, 20 µm.

**Supplementary Figure 4:**
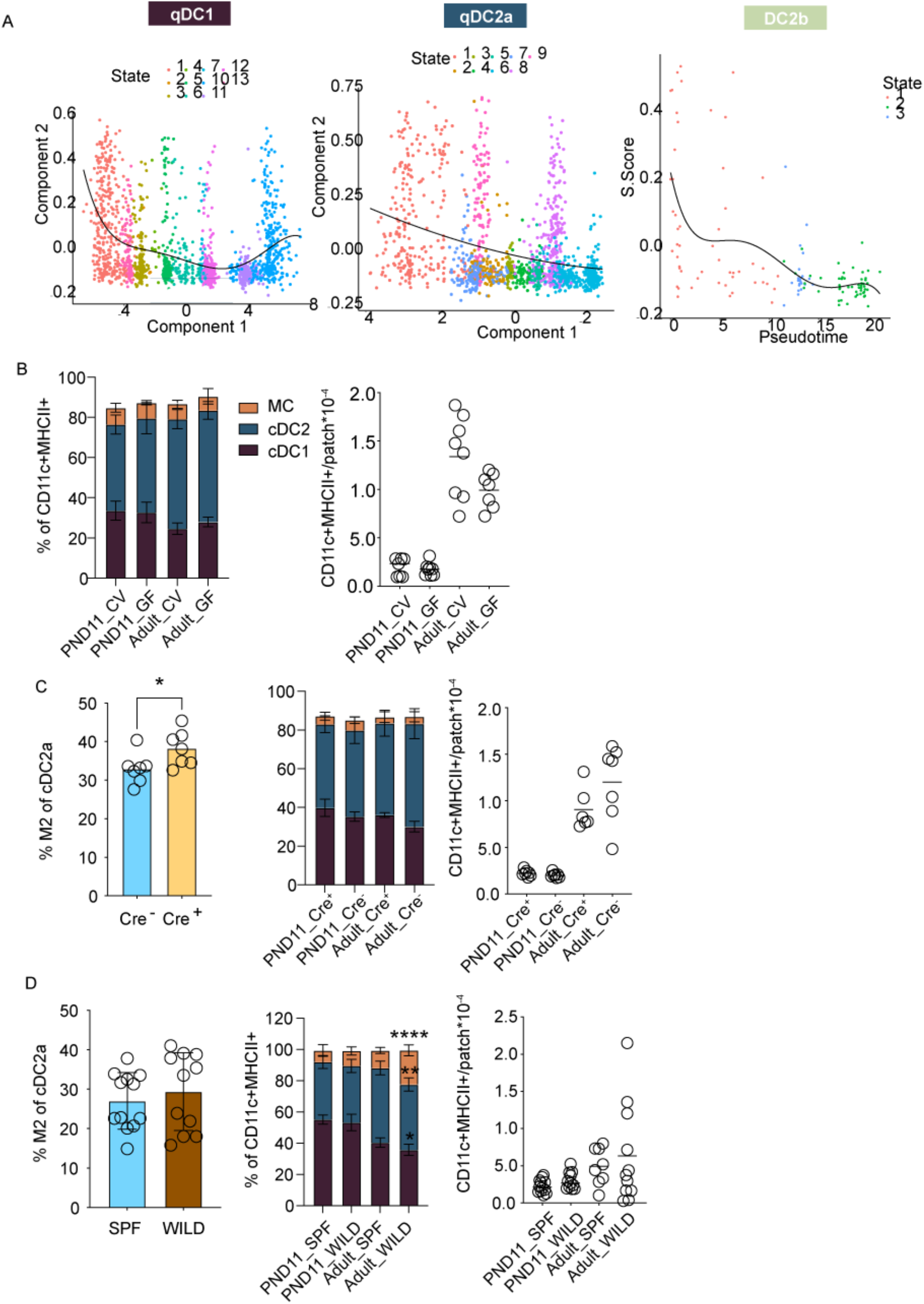
Maturation delay of PP cDC subsets during the postnatal period. (A) S-score projection onto the component 1 of the pseudotime trajectory for qDC1 (left panel), qDC2a (middle panel), and DC2b (right panel). (B) PP MNP subset composition (left panel) and absolute number of MNP per PP (right panel) in PND11 and adult germ-free (GF) or conventional (CV) mice determined by FACS (n=7-8, mean+SD and median). (C and D) Percentage of M2 (CD11b^+^) cells among cDC2 (Mann-Whitney U test, left panel); PP MNP subset composition (mean+SD, 2-way ANOVA, Tukey posttest, significance shown within age group and cell subset, middle panel) and absolute number of MNP per patch (median, Kruskall Wallis test, right panel) in (C) MyD88^fl/fl^ CD11Cre^+/-^ or Cre^-/-^ PND11 and adult mice (n=6-7) and (D) wildling (wild) and SPF, PND11 and adult mice (n=11-12). *, p<0.05; **, p<0.01; ***, p<0.001.

**Supplementary Figure 6:**
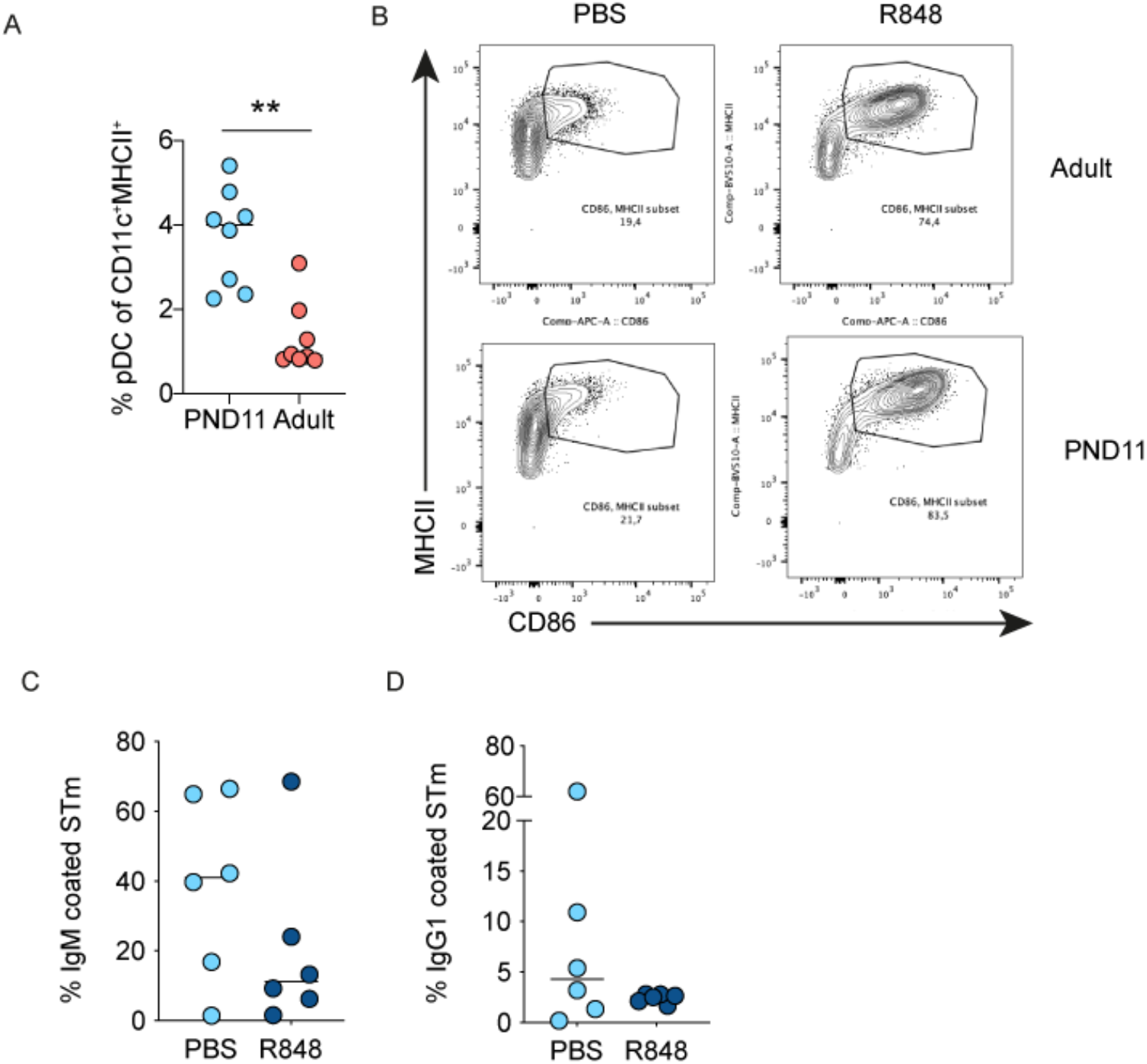
IFN I induces DC maturation in neonatal PP and modifies the adaptive immune response. **(A)** Percentage of BST2^hi^SIRPa^-^MHCII^lo^CD11c^+^CD45^+^live pDC among MNP in PP of PND11 and adult mice (n=8, mean, Mann Whitney U test). **(B)** Representative FACS plots showing the percentage of CD86^+^MHCII^+^ cells among MNP isolated from PP of PND11 (neonate) and adult mice 8h after oral gavage of 0.4 mg/g b.w. R848 or PBS. Percentage of **(C)** IgM^+^ and **(D)** IgG1^+^ *Salmonella* after incubation with sera from adult animals vaccinated against *Salmonella*±R848 as neonates (n=6, mena, Mann-Whitney U test). **, p<0.01.

## References

Abdel-Gadir, A., Stephen-Victor, E., Gerber, G.K., Noval Rivas, M., Wang, S., Harb, H., Wang, L., Li, N., Crestani, E., Spielman, S., et al. (2019). Microbiota therapy acts via a regulatory T cell MyD88/RORgammat pathway to suppress food allergy. Nat Med 25, 1164–1174.

Abt, M.C., Osborne, L.C., Monticelli, L.A., Doering, T.A., Alenghat, T., Sonnenberg, G.F., Paley, M.A., Antenus, M., Williams, K.L., Erikson, J., et al. (2012). Commensal bacteria calibrate the activation threshold of innate antiviral immunity. Immunity 37, 158–170.

Aramaki, T., Blanc-Mathieu, R., Endo, H., Ohkubo, K., Kanehisa, M., Goto, S., and Ogata, H. (2020). KofamKOALA: KEGG Ortholog assignment based on profile HMM and adaptive score threshold. Bioinformatics 36, 2251–2252.

Asnicar, F., Thomas, A.M., Beghini, F., Mengoni, C., Manara, S., Manghi, P., Zhu, Q., Bolzan, M., Cumbo, F., May, U., et al. (2020). Precise phylogenetic analysis of microbial isolates and genomes from metagenomes using PhyloPhlAn 3.0. Nat Commun 11, 2500.

Ayala, A.V.M. K.,; Hsu, C.-Y.; Terrazas, M. C.; Chu, H. (2021). Commensal bacteria promote type I interferon signaling to maintain immune tolerance.

Beura, L.K., Hamilton, S.E., Bi, K., Schenkel, J.M., Odumade, O.A., Casey, K.A., Thompson, E.A., Fraser, K.A., Rosato, P.C., Filali-Mouhim, A., et al. (2016). Normalizing the environment recapitulates adult human immune traits in laboratory mice. Nature 532, 512–516.

Bolger, A.M., Lohse, M., and Usadel, B. (2014). Trimmomatic: a flexible trimmer for Illumina sequence data. Bioinformatics 30, 2114–2120.

Bonnardel, J., Da Silva, C., Wagner, C., Bonifay, R., Chasson, L., Masse, M., Pollet, E., Dalod, M., Gorvel, J.P., and Lelouard, H. (2017). Distribution, location, and transcriptional profile of Peyer’s patch conventional DC subsets at steady state and under TLR7 ligand stimulation. Mucosal Immunol 10, 1412–1430.

Brown, C.C., Gudjonson, H., Pritykin, Y., Deep, D., Lavallee, V.P., Mendoza, A., Fromme, R., Mazutis, L., Ariyan, C., Leslie, C., et al. (2019). Transcriptional Basis of Mouse and Human Dendritic Cell Heterogeneity. Cell 179, 846–863 e824.

Bunker, J.J., Flynn, T.M., Koval, J.C., Shaw, D.G., Meisel, M., McDonald, B.D., Ishizuka, I.E., Dent, A.L., Wilson, P.C., Jabri, B., et al. (2015). Innate and Adaptive Humoral Responses Coat Distinct Commensal Bacteria with Immunoglobulin A. Immunity 43, 541–553.

Bushnell, B. (2014). BBMap: A Fast, Accurate, Splice-Aware Aligner.. 9th Annual Genomics of Energy & Environment Meeting.

Butler, A., Hoffman, P., Smibert, P., Papalexi, E., and Satija, R. (2018). Integrating single-cell transcriptomic data across different conditions, technologies, and species. Nat Biotechnol 36, 411–420.

Cabeza-Cabrerizo, M., Cardoso, A., Minutti, C.M., Pereira da Costa, M., and Reis, E.S.C. (2021). Dendritic Cells Revisited. Annu Rev Immunol 39, 131–166.

Cahenzli, J., Koller, Y., Wyss, M., Geuking, M.B., and McCoy, K.D. (2013). Intestinal microbial diversity during early-life colonization shapes long-term IgE levels. Cell Host Microbe 14, 559–570.

Constantinides, M.G., Link, V.M., Tamoutounour, S., Wong, A.C., Perez-Chaparro, P.J., Han, S.J., Chen, Y.E., Li, K., Farhat, S., Weckel, A., et al. (2019). MAIT cells are imprinted by the microbiota in early life and promote tissue repair. Science 366.

Costello, A., and Naimy, Z. (2019). Maternal, newborn, child and adolescent health: challenges for the next decade. Int Health 11, 349–352.

Countdown to 2030 Collaboration (2018). Countdown to 2030: tracking progress towards universal coverage for reproductive, maternal, newborn, and child health. Lancet 391, 1538–1548.

Da Silva, C., Wagner, C., Bonnardel, J., Gorvel, J.P., and Lelouard, H. (2017). The Peyer’s Patch Mononuclear Phagocyte System at Steady State and during Infection. Front Immunol 8, 1254.

Date, I., Koya, T., Sakamoto, T., Togi, M., Kawaguchi, H., Watanabe, A., Kato, T., Jr., and Shimodaira, S. (2020). Interferon-alpha-Induced Dendritic Cells Generated with Human Platelet Lysate Exhibit Elevated Antigen Presenting Ability to Cytotoxic T Lymphocytes. Vaccines (Basel) 9.

De Giovanni, M., Cutillo, V., Giladi, A., Sala, E., Maganuco, C.G., Medaglia, C., Di Lucia, P., Bono, E., Cristofani, C., Consolo, E., et al. (2020). Spatiotemporal regulation of type I interferon expression determines the antiviral polarization of CD4(+) T cells. Nat Immunol 21, 321–330.

Depner, M., Taft, D.H., Kirjavainen, P.V., Kalanetra, K.M., Karvonen, A.M., Peschel, S., Schmausser-Hechfellner, E., Roduit, C., Frei, R., Lauener, R., et al. (2020). Maturation of the gut microbiome during the first year of life contributes to the protective farm effect on childhood asthma. Nat Med 26, 1766–1775.

Ege, M.J., Mayer, M., Normand, A.C., Genuneit, J., Cookson, W.O., Braun-Fahrlander, C., Heederik, D., Piarroux, R., von Mutius, E., and Group, G.T.S. (2011). Exposure to environmental microorganisms and childhood asthma. N Engl J Med 364, 701–709.

Fiege, J.K., Block, K.E., Pierson, M.J., Nanda, H., Shepherd, F.K., Mickelson, C.K., Stolley, J.M., Matchett, W.E., Wijeyesinghe, S., Meyerholz, D.K., et al. (2021). Mice with diverse microbial exposure histories as a model for preclinical vaccine testing. Cell Host Microbe 29, 1815–1827 e1816.

Forster, R., Davalos-Misslitz, A.C., and Rot, A. (2008). CCR7 and its ligands: balancing immunity and tolerance. Nat Rev Immunol 8, 362–371.

G. B. D. Child Mortality Collaborators (2016). Global, regional, national, and selected subnational levels of stillbirths, neonatal, infant, and under-5 mortality, 1980-2015: a systematic analysis for the Global Burden of Disease Study 2015. Lancet 388, 1725–1774.

Gaboriau-Routhiau, V., Rakotobe, S., Lecuyer, E., Mulder, I., Lan, A., Bridonneau, C., Rochet, V., Pisi, A., De Paepe, M., Brandi, G., et al. (2009). The key role of segmented filamentous bacteria in the coordinated maturation of gut helper T cell responses. Immunity 31, 677–689.

Ganal, S.C., Sanos, S.L., Kallfass, C., Oberle, K., Johner, C., Kirschning, C., Lienenklaus, S., Weiss, S., Staeheli, P., Aichele, P., et al. (2012). Priming of natural killer cells by nonmucosal mononuclear phagocytes requires instructive signals from commensal microbiota. Immunity 37, 171–186.

Ganeshan, K., Nikkanen, J., Man, K., Leong, Y.A., Sogawa, Y., Maschek, J.A., Van Ry, T., Chagwedera, D.N., Cox, J.E., and Chawla, A. (2019). Energetic Trade-Offs and Hypometabolic States Promote Disease Tolerance. Cell 177, 399–413 e312.

Gardner, J.M., Devoss, J.J., Friedman, R.S., Wong, D.J., Tan, Y.X., Zhou, X., Johannes, K.P., Su, M.A., Chang, H.Y., Krummel, M.F., et al. (2008). Deletional tolerance mediated by extrathymic Aire-expressing cells. Science 321, 843–847.

Gavin, M.A., and Bevan, M.J. (1995). Increased peptide promiscuity provides a rationale for the lack of N regions in the neonatal T cell repertoire. Immunity 3, 793–800.

Godon, J.J., Zumstein, E., Dabert, P., Habouzit, F., and Moletta, R. (1997). Molecular microbial diversity of an anaerobic digestor as determined by small-subunit rDNA sequence analysis. Appl Environ Microbiol 63, 2802–2813.

Gollwitzer, E.S., Saglani, S., Trompette, A., Yadava, K., Sherburn, R., McCoy, K.D., Nicod, L.P., Lloyd, C.M., and Marsland, B.J. (2014). Lung microbiota promotes tolerance to allergens in neonates via PD-L1. Nat Med 20, 642–647.

Gough, D.J., Messina, N.L., Clarke, C.J., Johnstone, R.W., and Levy, D.E. (2012). Constitutive type I interferon modulates homeostatic balance through tonic signaling. Immunity 36, 166–174.

Graham, D.B., Luo, C., O’Connell, D.J., Lefkovith, A., Brown, E.M., Yassour, M., Varma, M., Abelin, J.G., Conway, K.L., Jasso, G.J., et al. (2018). Antigen discovery and specification of immunodominance hierarchies for MHCII-restricted epitopes. Nat Med 24, 1762–1772.

Harbeson, D., Francis, F., Bao, W., Amenyogbe, N.A., and Kollmann, T.R. (2018). Energy Demands of Early Life Drive a Disease Tolerant Phenotype and Dictate Outcome in Neonatal Bacterial Sepsis. Front Immunol 9, 1918.

Henrick, B.M., Rodriguez, L., Lakshmikanth, T., Pou, C., Henckel, E., Arzoomand, A., Olin, A., Wang, J., Mikes, J., Tan, Z., et al. (2021). Bifidobacteria-mediated immune system imprinting early in life. Cell.

Herzenberg, L.A., and Herzenberg, L.A. (1989). Toward a layered immune system. Cell 59, 953–954.

Hooper, L.V., Littman, D.R., and Macpherson, A.J. (2012). Interactions between the microbiota and the immune system. Science 336, 1268–1273.

Hornef, M.W., and Torow, N. (2020). ‘Layered immunity’ and the ‘neonatal window of opportunity’ - timed succession of non-redundant phases to establish mucosal host-microbial homeostasis after birth. Immunology 159, 15–25.

Jego, G., Palucka, A.K., Blanck, J.P., Chalouni, C., Pascual, V., and Banchereau, J. (2003). Plasmacytoid dendritic cells induce plasma cell differentiation through type I interferon and interleukin 6. Immunity 19, 225–234.

Jiang, H.Q., Bos, N.A., and Cebra, J.J. (2001). Timing, localization, and persistence of colonization by segmented filamentous bacteria in the neonatal mouse gut depend on immune status of mothers and pups. Infect Immun 69, 3611–3617.

Kanaya, T., Hase, K., Takahashi, D., Fukuda, S., Hoshino, K., Sasaki, I., Hemmi, H., Knoop, K.A., Kumar, N., Sato, M., et al. (2012). The Ets transcription factor Spi-B is essential for the differentiation of intestinal microfold cells. Nat Immunol 13, 729–736.

Kawashima, T., Ikari, N., Watanabe, Y., Kubota, Y., Yoshio, S., Kanto, T., Motohashi, S., Shimojo, N., and Tsuji, N.M. (2018). Double-Stranded RNA Derived from Lactic Acid Bacteria Augments Th1 Immunity via Interferon-beta from Human Dendritic Cells. Front Immunol 9, 27.

Kawashima, T., Kosaka, A., Yan, H., Guo, Z., Uchiyama, R., Fukui, R., Kaneko, D., Kumagai, Y., You, D.J., Carreras, J., et al. (2013). Double-stranded RNA of intestinal commensal but not pathogenic bacteria triggers production of protective interferon-beta. Immunity 38, 1187–1197.

Kirjavainen, P.V., Karvonen, A.M., Adams, R.I., Taubel, M., Roponen, M., Tuoresmaki, P., Loss, G., Jayaprakash, B., Depner, M., Ege, M.J., et al. (2019). Farm-like indoor microbiota in non-farm homes protects children from asthma development. Nat Med 25, 1089–1095.

Klindworth, A., Pruesse, E., Schweer, T., Peplies, J., Quast, C., Horn, M., and Glockner, F.O. (2013). Evaluation of general 16S ribosomal RNA gene PCR primers for classical and next-generation sequencing-based diversity studies. Nucleic Acids Res 41, e1.

Koch, M.A., Reiner, G.L., Lugo, K.A., Kreuk, L.S., Stanbery, A.G., Ansaldo, E., Seher, T.D., Ludington, W.B., and Barton, G.M. (2016). Maternal IgG and IgA Antibodies Dampen Mucosal T Helper Cell Responses in Early Life. Cell 165, 827–841.

Kollmann, T.R., Marchant, A., and Way, S.S. (2020). Vaccination strategies to enhance immunity in neonates. Science 368, 612–615.

Kotloff, K.L., Nataro, J.P., Blackwelder, W.C., Nasrin, D., Farag, T.H., Panchalingam, S., Wu, Y., Sow, S.O., Sur, D., Breiman, R.F., et al. (2013). Burden and aetiology of diarrhoeal disease in infants and young children in developing countries (the Global Enteric Multicenter Study, GEMS): a prospective, case-control study. Lancet 382, 209–222.

Koya, T., Yanagisawa, R., Higuchi, Y., Sano, K., and Shimodaira, S. (2017). Interferon-alpha-inducible Dendritic Cells Matured with OK-432 Exhibit TRAIL and Fas Ligand Pathway-mediated Killer Activity. Sci Rep 7, 42145.

Kuma, A., Hatano, M., Matsui, M., Yamamoto, A., Nakaya, H., Yoshimori, T., Ohsumi, Y., Tokuhisa, T., and Mizushima, N. (2004). The role of autophagy during the early neonatal starvation period. Nature 432, 1032–1036.

Le Bon, A., Schiavoni, G., D’Agostino, G., Gresser, I., Belardelli, F., and Tough, D.F. (2001). Type i interferons potently enhance humoral immunity and can promote isotype switching by stimulating dendritic cells in vivo. Immunity 14, 461–470.

Lelouard, H., Fallet, M., de Bovis, B., Meresse, S., and Gorvel, J.P. (2012). Peyer’s patch dendritic cells sample antigens by extending dendrites through M cell-specific transcellular pores. Gastroenterology 142, 592–601 e593.

Lelouard, H., Mailfert, S., and Fallet, M. (2018). A ten-color spectral imagingstrategy to reveal localization of gut immune cell subsets.

Lesker, T.R., Durairaj, A.C., Galvez, E.J.C., Lagkouvardos, I., Baines, J.F., Clavel, T., Sczyrba, A., McHardy, A.C., and Strowig, T. (2020). An Integrated Metagenome Catalog Reveals New Insights into the Murine Gut Microbiome. Cell Rep 30, 2909–2922 e2906.

Letunic, I., and Bork, P. (2019). Interactive Tree Of Life (iTOL) v4: recent updates and new developments. Nucleic Acids Res 47, W256–W259.

Levine, M.M., Nasrin, D., Acacio, S., Bassat, Q., Powell, H., Tennant, S.M., Sow, S.O., Sur, D., Zaidi, A.K.M., Faruque, A.S.G., et al. (2020). Diarrhoeal disease and subsequent risk of death in infants and children residing in low-income and middle-income countries: analysis of the GEMS case-control study and 12-month GEMS-1A follow-on study. Lancet Glob Health 8, e204–e214.

Li, D., Liu, C.M., Luo, R., Sadakane, K., and Lam, T.W. (2015). MEGAHIT: an ultra-fast single-node solution for large and complex metagenomics assembly via succinct de Bruijn graph. Bioinformatics 31, 1674–1676.

Li, N., van Unen, V., Abdelaal, T., Guo, N., Kasatskaya, S.A., Ladell, K., McLaren, J.E., Egorov, E.S., Izraelson, M., Chuva de Sousa Lopes, S.M., et al. (2019). Memory CD4(+) T cells are generated in the human fetal intestine. Nat Immunol 20, 301–312.

Lienenklaus, S., Cornitescu, M., Zietara, N., Lyszkiewicz, M., Gekara, N., Jablonska, J., Edenhofer, F., Rajewsky, K., Bruder, D., Hafner, M., et al. (2009). Novel reporter mouse reveals constitutive and inflammatory expression of IFN-beta in vivo. J Immunol 183, 3229–3236.

Longhi, M.P., Trumpfheller, C., Idoyaga, J., Caskey, M., Matos, I., Kluger, C., Salazar, A.M., Colonna, M., and Steinman, R.M. (2009). Dendritic cells require a systemic type I interferon response to mature and induce CD4+ Th1 immunity with poly IC as adjuvant. J Exp Med 206, 1589–1602.

Luciani, C., Hager, F.T., Cerovic, V., and Lelouard, H. (2021). Dendritic cell functions in the inductive and effector sites of intestinal immunity. Mucosal Immunol.

Lynn, M.A., Tumes, D.J., Choo, J.M., Sribnaia, A., Blake, S.J., Leong, L.E.X., Young, G.P., Marshall, H.S., Wesselingh, S.L., Rogers, G.B., et al. (2018). Early-Life Antibiotic-Driven Dysbiosis Leads to Dysregulated Vaccine Immune Responses in Mice. Cell Host Microbe 23, 653–660 e655.

MacLennan, C.A., and Saul, A. (2014). Vaccines against poverty. Proc Natl Acad Sci U S A 111, 12307–12312.

Marie, I.J., Brambilla, L., Azzouz, D., Chen, Z., Baracho, G.V., Arnett, A., Li, H.S., Liu, W., Cimmino, L., Chattopadhyay, P., et al. (2021). Tonic interferon restricts pathogenic IL-17-driven inflammatory disease via balancing the microbiome. Elife 10.

Martinez, I., Maldonado-Gomez, M.X., Gomes-Neto, J.C., Kittana, H., Ding, H., Schmaltz, R., Joglekar, P., Cardona, R.J., Marsteller, N.L., Kembel, S.W., et al. (2018). Experimental evaluation of the importance of colonization history in early-life gut microbiota assembly. Elife 7.

Mathers, C., Stevens, G., Hogan, D., Mahanani, W.R., and Ho, J. (2017). Global and Regional Causes of Death: Patterns and Trends, 2000-15. In Disease Control Priorities: Improving Health and Reducing Poverty, rd, D.T. Jamison, H. Gelband, S. Horton, P. Jha, R. Laxminarayan, C.N. Mock, and R. Nugent, eds. (Washington (DC)).

Medzhitov, R., Schneider, D.S., and Soares, M.P. (2012). Disease tolerance as a defense strategy. Science 335, 936–941.

Mesev, E.V., LeDesma, R.A., and Ploss, A. (2019). Decoding type I and III interferon signalling during viral infection. Nat Microbiol 4, 914–924.

Mishra, A., Lai, G.C., Yao, L.J., Aung, T.T., Shental, N., Rotter-Maskowitz, A., Shepherdson, E., Singh, G.S.N., Pai, R., Shanti, A., et al. (2021). Microbial exposure during early human development primes fetal immune cells. Cell 184, 3394–3409 e3320.

Moor, K., Wotzka, S.Y., Toska, A., Diard, M., Hapfelmeier, S., and Slack, E. (2016). Peracetic Acid Treatment Generates Potent Inactivated Oral Vaccines from a Broad Range of Culturable Bacterial Species. Front Immunol 7, 34.

Olszak, T., An, D., Zeissig, S., Vera, M.P., Richter, J., Franke, A., Glickman, J.N., Siebert, R., Baron, R.M., Kasper, D.L., et al. (2012). Microbial exposure during early life has persistent effects on natural killer T cell function. Science 336, 489–493.

Oppong, T.B., Yang, H., Amponsem-Boateng, C., Kyere, E.K.D., Abdulai, T., Duan, G., and Opolot, G. (2020). Enteric pathogens associated with gastroenteritis among children under 5 years in sub-Saharan Africa: a systematic review and meta-analysis. Epidemiol Infect 148, e64.

Ozkul, C., Ruiz, V.E., Battaglia, T., Xu, J., Roubaud-Baudron, C., Cadwell, K., Perez-Perez, G.I., and Blaser, M.J. (2020). A single early-in-life antibiotic course increases susceptibility to DSS-induced colitis. Genome Med 12, 65.

Pabst, O., and Slack, E. (2020). IgA and the intestinal microbiota: the importance of being specific. Mucosal Immunol 13, 12–21.

Pantoja-Feliciano, I.G., Clemente, J.C., Costello, E.K., Perez, M.E., Blaser, M.J., Knight, R., and Dominguez-Bello, M.G. (2013). Biphasic assembly of the murine intestinal microbiota during early development. ISME J 7, 1112–1115.

Papaioannou, N.E., Salei, N., Rambichler, S., Ravi, K., Popovic, J., Kuntzel, V., Lehmann, C.H.K., Fiancette, R., Salvermoser, J., Gajdasik, D.W., et al. (2021). Environmental signals rather than layered ontogeny imprint the function of type 2 conventional dendritic cells in young and adult mice. Nat Commun 12, 464.

Prudden, H.J., Hasso-Agopsowicz, M., Black, R.E., Troeger, C., Reiner, R.C., Breiman, R.F., Jit, M., Kang, G., Lamberti, L., Lanata, C.F., et al. (2020). Meeting Report: WHO Workshop on modelling global mortality and aetiology estimates of enteric pathogens in children under five. Cape Town, 28-29th November 2018. Vaccine 38, 4792–4800.

Rivera, C.A., Randrian, V., Richer, W., Gerber-Ferder, Y., Delgado, M.G., Chikina, A.S., Frede, A., Sorini, C., Maurin, M., Kammoun-Chaari, H., et al. (2021). Epithelial colonization by gut dendritic cells promotes their functional diversification. Immunity.

Rosshart, S.P., Herz, J., Vassallo, B.G., Hunter, A., Wall, M.K., Badger, J.H., McCulloch, J.A., Anastasakis, D.G., Sarshad, A.A., Leonardi, I., et al. (2019). Laboratory mice born to wild mice have natural microbiota and model human immune responses. Science 365.

Rosshart, S.P., Vassallo, B.G., Angeletti, D., Hutchinson, D.S., Morgan, A.P., Takeda, K., Hickman, H.D., McCulloch, J.A., Badger, J.H., Ajami, N.J., et al. (2017). Wild Mouse Gut Microbiota Promotes Host Fitness and Improves Disease Resistance. Cell 171, 1015–1028 e1013.

Roswall, J., Olsson, L.M., Kovatcheva-Datchary, P., Nilsson, S., Tremaroli, V., Simon, M.C., Kiilerich, P., Akrami, R., Kramer, M., Uhlen, M., et al. (2021). Developmental trajectory of the healthy human gut microbiota during the first 5 years of life. Cell Host Microbe 29, 765–776 e763.

Salonen, A., Nikkila, J., Jalanka-Tuovinen, J., Immonen, O., Rajilic-Stojanovic, M., Kekkonen, R.A., Palva, A., and de Vos, W.M. (2010). Comparative analysis of fecal DNA extraction methods with phylogenetic microarray: effective recovery of bacterial and archaeal DNA using mechanical cell lysis. J Microbiol Methods 81, 127–134.

Satija, R., Farrell, J.A., Gennert, D., Schier, A.F., and Regev, A. (2015). Spatial reconstruction of single-cell gene expression data. Nat Biotechnol 33, 495–502.

Satopa, V.A. J.,; Irwin, D.; Raghavan, B. (2011). Finding a “Kneedle” in a Haystack: Detecting Knee Points in System Behavior. Paper presented at: 31st International Conference on Distributed Computing Systems Workshops.

Schaupp, L., Muth, S., Rogell, L., Kofoed-Branzk, M., Melchior, F., Lienenklaus, S., Ganal-Vonarburg, S.C., Klein, M., Guendel, F., Hain, T., et al. (2020). Microbiota-Induced Type I Interferons Instruct a Poised Basal State of Dendritic Cells. Cell 181, 1080–1096 e1019.

Schreurs, R., Baumdick, M.E., Sagebiel, A.F., Kaufmann, M., Mokry, M., Klarenbeek, P.L., Schaltenberg, N., Steinert, F.L., van Rijn, J.M., Drewniak, A., et al. (2019). Human Fetal TNF-alpha-Cytokine-Producing CD4(+) Effector Memory T Cells Promote Intestinal Development and Mediate Inflammation Early in Life. Immunity 50, 462–476 e468.

Schuijs, M.J., Willart, M.A., Vergote, K., Gras, D., Deswarte, K., Ege, M.J., Madeira, F.B., Beyaert, R., van Loo, G., Bracher, F., et al. (2015). Farm dust and endotoxin protect against allergy through A20 induction in lung epithelial cells. Science 349, 1106–1110.

Seemann, T. (2014). Prokka: rapid prokaryotic genome annotation. Bioinformatics 30, 2068–2069.

Spadaro, F., Lapenta, C., Donati, S., Abalsamo, L., Barnaba, V., Belardelli, F., Santini, S.M., and Ferrantini, M. (2012). IFN-alpha enhances cross-presentation in human dendritic cells by modulating antigen survival, endocytic routing, and processing. Blood 119, 1407–1417.

Stebegg, M., Bignon, A., Hill, D.L., Silva-Cayetano, A., Krueger, C., Vanderleyden, I., Innocentin, S., Boon, L., Wang, J., Zand, M.S., et al. (2020). Rejuvenating conventional dendritic cells and T follicular helper cell formation after vaccination. Elife 9.

Stefan, K.L., Kim, M.V., Iwasaki, A., and Kasper, D.L. (2020). Commensal Microbiota Modulation of Natural Resistance to Virus Infection. Cell 183, 1312–1324 e1310.

Taniguchi, T., and Takaoka, A. (2001). A weak signal for strong responses: interferon-alpha/beta revisited. Nat Rev Mol Cell Biol 2, 378–386.

Torow, N., Dittrich-Breiholz, O., and Hornef, M.W. (2015a). Transcriptional profiling of intestinal CD4(+) T cells in the neonatal and adult mice. Genom Data 5, 371–374.

Torow, N., Yu, K., Hassani, K., Freitag, J., Schulz, O., Basic, M., Brennecke, A., Sparwasser, T., Wagner, N., Bleich, A., et al. (2015b). Active suppression of intestinal CD4(+)TCRalphabeta(+) T-lymphocyte maturation during the postnatal period. Nat Commun 6, 7725.

Trapnell, C., Cacchiarelli, D., Grimsby, J., Pokharel, P., Li, S., Morse, M., Lennon, N.J., Livak, K.J., Mikkelsen, T.S., and Rinn, J.L. (2014). The dynamics and regulators of cell fate decisions are revealed by pseudotemporal ordering of single cells. Nat Biotechnol 32, 381–386.

van Best, N., Rolle-Kampczyk, U., Schaap, F.G., Basic, M., Olde Damink, S.W.M., Bleich, A., Savelkoul, P.H.M., von Bergen, M., Penders, J., and Hornef, M.W. (2020). Bile acids drive the newborn’s gut microbiota maturation. Nat Commun 11, 3692.

Vatanen, T., Kostic, A.D., d’Hennezel, E., Siljander, H., Franzosa, E.A., Yassour, M., Kolde, R., Vlamakis, H., Arthur, T.D., Hamalainen, A.M., et al. (2016a). Variation in Microbiome LPS Immunogenicity Contributes to Autoimmunity in Humans. Cell 165, 1551.

Vatanen, T., Kostic, A.D., d’Hennezel, E., Siljander, H., Franzosa, E.A., Yassour, M., Kolde, R., Vlamakis, H., Arthur, T.D., Hamalainen, A.M., et al. (2016b). Variation in Microbiome LPS Immunogenicity Contributes to Autoimmunity in Humans. Cell 165, 842–853.

Wagner, C., Bonnardel, J., Da Silva, C., Spinelli, L., Portilla, C.A., Tomas, J., Lagier, M., Chasson, L., Masse, M., Dalod, M., et al. (2020). Differentiation Paths of Peyer’s Patch LysoDCs Are Linked to Sampling Site Positioning, Migration, and T Cell Priming. Cell Rep 31, 107479.

Wang, J., Lareau, C.A., Bautista, J.L., Gupta, A.R., Sandor, K., Germino, J., Yin, Y., Arvedson, M.P., Reeder, G.C., Cramer, N.T., et al. (2021). Single-cell multiomics defines tolerogenic extrathymic Aire-expressing populations with unique homology to thymic epithelium. Sci Immunol 6, eabl5053.

Weinstein, P.D., and Cebra, J.J. (1991). The preference for switching to IgA expression by Peyer’s patch germinal center B cells is likely due to the intrinsic influence of their microenvironment. J Immunol 147, 4126–4135.

Yang, X.L., Wang, G., Xie, J.Y., Li, H., Chen, S.X., Liu, W., and Zhu, S.J. (2021). The Intestinal Microbiome Primes Host Innate Immunity against Enteric Virus Systemic Infection through Type I Interferon. mBio 12.

Yatsunenko, T., Rey, F.E., Manary, M.J., Trehan, I., Dominguez-Bello, M.G., Contreras, M., Magris, M., Hidalgo, G., Baldassano, R.N., Anokhin, A.P., et al. (2012). Human gut microbiome viewed across age and geography. Nature 486, 222–227.

Yrlid, U., Milling, S.W., Miller, J.L., Cartland, S., Jenkins, C.D., and MacPherson, G.G. (2006). Regulation of intestinal dendritic cell migration and activation by plasmacytoid dendritic cells, TNF-alpha and type 1 IFNs after feeding a TLR7/8 ligand. J Immunol 176, 5205–5212.

Zanvit, P., Konkel, J.E., Jiao, X., Kasagi, S., Zhang, D., Wu, R., Chia, C., Ajami, N.J., Smith, D.P., Petrosino, J.F., et al. (2015). Antibiotics in neonatal life increase murine susceptibility to experimental psoriasis. Nat Commun 6, 8424.

Zhang, K., Dupont, A., Torow, N., Gohde, F., Leschner, S., Lienenklaus, S., Weiss, S., Brinkmann, M.M., Kuhnel, M., Hensel, M., et al. (2014). Age-dependent enterocyte invasion and microcolony formation by Salmonella. PLoS Pathog 10, e1004385.

Zietara, N., Lyszkiewicz, M., Gekara, N., Puchalka, J., Dos Santos, V.A., Hunt, C.R., Pandita, T.K., Lienenklaus, S., and Weiss, S. (2009). Absence of IFN-beta impairs antigen presentation capacity of splenic dendritic cells via down-regulation of heat shock protein 70. J Immunol 183, 1099–1109.

